# Unsupervised learning reveals rapid gait adaption after leg loss and regrowth in spiders

**DOI:** 10.1101/2025.01.23.634080

**Authors:** Suzanne Amador Kane, Brooke L. Quinn, Xuanyi Kris Wu, Sarah Y. Xi, Michael F. Ochs, S. Tonia Hsieh

## Abstract

Many arthropods and some vertebrates can voluntarily lose (autotomize) limbs during antagonistic encounters, and some can regenerate functional replacements. Spiders in particular frequently autotomize one or more legs. In this study, we investigated the time course of locomotor recovery after leg loss and regeneration in juvenile tarantulas (Arachnida: Araneae) with no prior experience of autotomy. We recorded high-speed video of spiders running with all legs intact, then immediately after, and again one day after, they autotomized two legs. The legs were allowed to regenerate, and the same sequence of experiments repeated. Running performance, posture, and path tortuosity were measured from video tracking. Spiders were found to resume their pre-autotomy speed and stride frequency after leg regeneration and in ≤1 day after both autotomies; furthermore, path tortuosity was unaffected by these treatments. They adjusted their posture to compensate for missing legs, spreading their remaining legs and running with their bodies rotated 11-15 deg from their velocity. To analyze gaits, we applied unsupervised machine learning for the first time to measured kinematic data in combination with gait space metrics. Spiders were found to robustly adopt new gait patterns immediately after losing legs, with no evidence of learning. This novel clustering approach both demonstrated concordance with previously-hypothesized gaits and revealed transitions between and variations within these patterns. More generally, clustering in gait space enables the identification of patterns of leg motions in large datasets that correspond to either known gaits or undiscovered behaviors.

## Introduction

Autotomy, the voluntary loss of an appendage, is a common strategy used for predator evasion in the animal kingdom (Emberts et al., 2019; Fleming et al., 2007). Leg autotomy is especially common among spiders (Arachnida, order Araneae), with 5-40% of specimens observed in the field reported as missing one or more limbs (Brown et al., 2018; Brueseke et al., 2001; Johnson and Jakob, 1999; Wrinn and Uetz, 2007). If the loss of one or more limbs compromises a spider’s locomotion, this could impede its ability to capture prey and evade predators while navigating diverse, complex habitats. In practice, the loss of one or more legs was found to result in significantly reduced running speed for spiders in some (Apontes and Brown, 2005; Brown and Formanowicz, 2012; Gerald et al., 2017), but not all (Boehm et al., 2021; Brueseke et al., 2001; Wilshin et al., 2018) previous studies. This raises the question of whether and how these arachnids compensate for the loss of one or more limbs. Furthermore, while the juveniles of many species of spiders are able to regenerate functional legs after molting (Foelix, 2011), no previous studies have explored how leg regeneration influences locomotive performance in spiders, and whether spiders respond differently to a subsequent repeated appendage loss.

One might expect the loss of a leg used in a favored gait to especially impact an arachnid’s locomotion. In particular, arthropods often locomote using “trotting” gaits in which two sets of diagonally-opposed alternating limbs (“pods”) move in synchrony. Just as six-legged insects commonly run using an alternating tripod gait (Fig. 1A), eight-legged arachnids often run with an alternating tetrapod gait employing synchronously moving groups of four diagonally-opposed alternating limbs (tetrapods). (Fig. 1B, Movie 1) Gaits similar to this have been reported for spiders (Biancardi and Silva-Pereyra, 2020; Biancardi et al., 2011; Land, 1972; Silva-Pereyra et al., 2019; Spagna and Peattie, 2012; Weihmann, 2013; Weihmann, 2020; Wilson, 1967), scorpions (Bowerman, 1975a), pseudoscorpions (Tross et al., 2022), and mites (Weihmann et al., 2015). Therefore, the loss of two legs from a single tetrapod has been proposed as an especially destabilizing injury. Indeed, tarantulas (*Dugesiella hentzi* (Girard)), wolf spiders (Lycosidae, *Pardosa* sp. Koch 1847), and the scorpion *Hadrurus arizonensis* with one foreleg and the opposing hindleg from the same tetrapod autotomized have been found to adopt one of two gaits: 1) an “ablated tetrapod” gait, in which they alternate using the intact tetrapod and a bipod formed by two legs remaining from the ablated tetrapod (Fig. 1C, Movie 1); or 2) a “modified tripod” gait, in which they move the remaining six limbs in an adaptive version of the alternating tripod gait (Fig. 1D, Movie 1) (Bowerman, 1975b; Wilshin et al., 2018; Wilson, 1967). These altered gaits have different trade-offs: adjusting to the modified tripod gait requires altering the phasing of leg motions, while using the ablated tetrapod gait requires moving with only two legs on the ground for part of each stride.

**Fig. 1.**
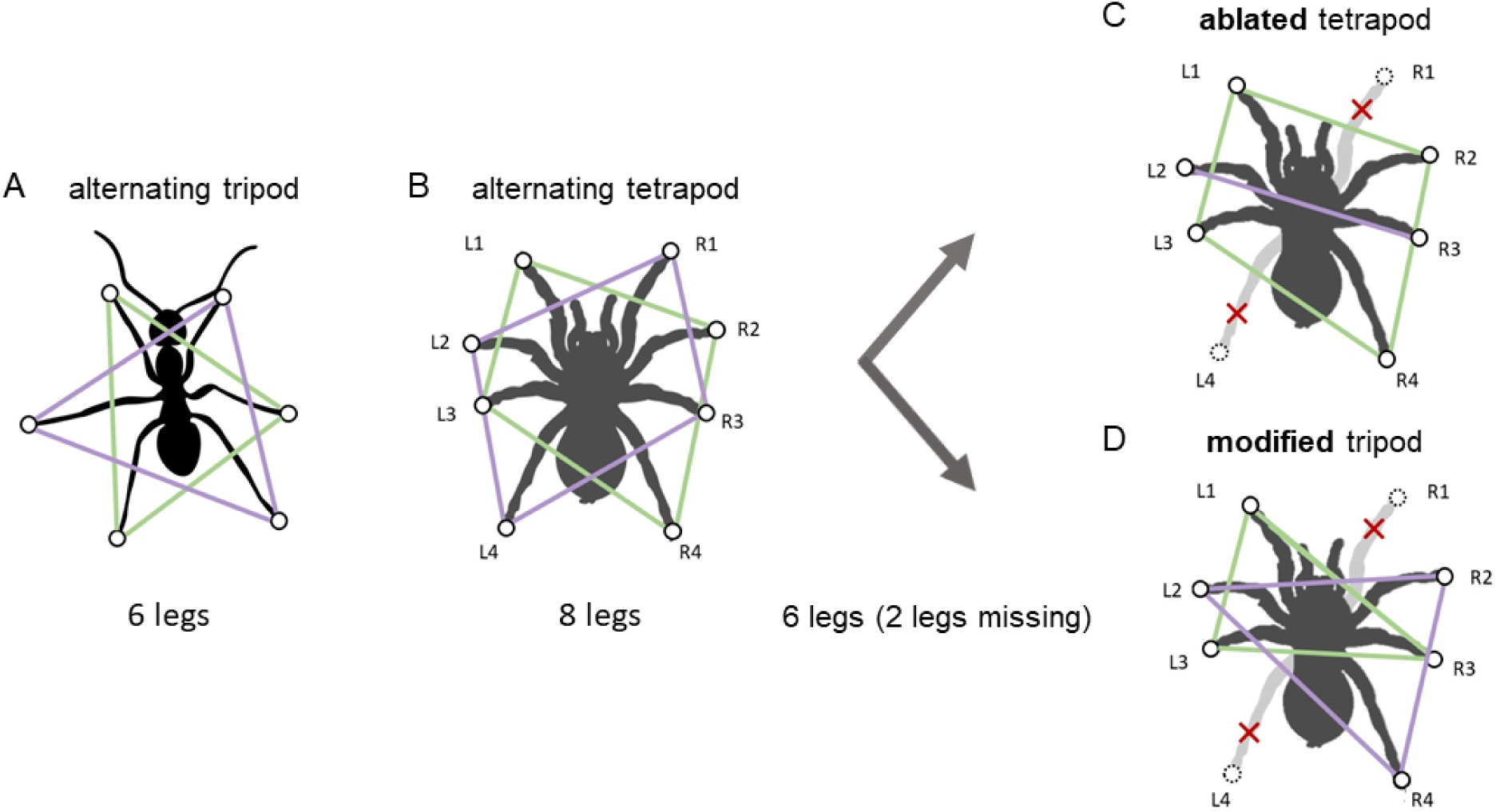
Illustration of typical gaits used by A) hexapods (e.g., 6-legged insects) and B) octopods (e.g., 8-legged spiders). The two sets of legs moved synchronously (tripods of 3 legs for hexapods and tetrapods of 4 legs for octopods) are indicated by green and purple lines respectively. Gaits used by spiders that have lost 2 legs include C) the ablated tetrapod, in which one tetrapod continues the same motion as pre-autotomy (green lines) and a bipod consisting of the other two intact legs move together (purple line), and D) the modified tripod, in which the remaining intact six legs move in two alternating tripods, each of which incorporates legs from both original tetrapods. The labels used to denote spider legs in this study are indicated in B-D. (The ant silhouette in Fig. 1A is from https://commons.wikimedia.org/wiki/File:Fourmi02.svg#file by Meul, accessed May 2, 2024 with a <u>Creative Commons Attribution-Share Alike 3.0 Unported</u> license)

Various adaptations in response to leg autotomy have been reported for different arthropods. For example, many studies have reported that leg autotomy can result in reduced speed in a variety of taxa (Fleming et al., 2007) (Lutzy and Morse, 2008; Pfeiffenberger and Hsieh, 2021; Prestholdt et al., 2022) (Escalante and O’Brien, 2024) (O’Neil et al., 2024) (Han, 2024) (Meshkani et al., 2023). Arthropods that have been reported to adopt modified gait patterns after leg loss include the stinkbugs (Han, 2024), harvestmen (Opiliones: Sclerosomatidae), a group of 8-legged arachnids, and crabs (Barnes, 1975; Herreid and Full, 1986). By contrast, cockroaches that had lost either one or two legs continued to locomote using the same speed and the same motions of the remaining legs as observed for the alternating tripod gait (Jayaram, 2015; Saintsing, 2022). Some species also respond by modifying their posture and leg use post-autotomy: e.g., the harvestman *Prionostemma Pocock* adopts a more crouched posture and adapts legs ordinarily used for sensing to locomotion, while cockroaches and water striders widen the stance of their remaining legs (Jayaram, 2015; Saintsing, 2022) (Meshkani et al., 2023) and stinkbugs and ants alter the posture and swing range of their remaining legs to fill in in missing functional space due to lost legs (Han, 2024) (Zeng et al., 2024). A few arthropods have been reported to walk after leg loss using more tortuous (e.g., more nonlinear and sinuous) trajectories, including harvestmen (Escalante and O’Brien, 2024) and the water-walking insect *Microvelia* (O’Neil et al., 2024).

There have been relatively few investigations of the time course of recovery of locomotor performance after limb autotomy in arthropods. Harvestmen that lost 3 legs experienced an initial 40% speed decrease and a return to pre-autotomy values after 2 days (Escalante et al., 2020). Cockroaches used the same gaits modifications 3 days post-autotomy as immediately after (Hughes, 1957). Water-walking insects moved at reduced speed in a modified gait for an hour after losing 2 legs, but resumed the tripod gait at higher speed within 24 hours (O’Neil et al., 2024).

We are aware of only one study of the effect of limb regeneration post-autotomy on locomotion. The purple shore crab (*Hemigrapsus nudus*) was found to sprint at reduced top speeds following leg autotomy, but there was no difference in top sprint speed between controls and crabs with regenerated legs (Prestholdt et al., 2022).

Here we report on locomotion by spiders using high-speed 2D video of running recorded before and after experiencing two successive two-leg autotomies and regrowing both legs in between. To analyze these data, we tracked the spiders’ body and leg motions and used these results to compute measures of their locomotor performance.

Based on previous research, we tested a series of hypotheses. First, we expected that autotomized spiders would have reduced locomotor performance (i.e., reduced speed, lower stability, etc.) compared to intact specimens. To test this, we looked for changes in speed and body yaw, changes in the path tortuosity (a measure of the deviation from a straight path), variability in stride frequency and length, and changes in various proposed stability measures.

Second, we expected tarantulas to widen their stance and increase tarsal range of motion to enhance locomotor stability, as reported for cockroaches, water striders and stinkbugs (Han, 2024; Meshkani et al., 2023; Saintsing, 2022)

Third, we hypothesized that intact spiders would use the alternating tetrapod gait pre-autotomy and after leg regeneration, but switch to a combination of the ablated tetrapod and modified tripod gaits post-autotomy (Fig. 1B-D).

Previous studies of locomotion have classified gaits based on the differences in oscillation phase between various pairs of legs. For example, gait stepping patterns can be classified by specifying the subsets of tarsi that move as an approximately synchronous unit and those that alternate in their motion (Biancardi and Silva-Pereyra, 2020). Another approach is to compare the set of phase differences between the oscillation of pairs of legs that define a gait using a statistical test of significance suitable for circular statistics, such as the Kuiper two-sample test (Weihmann et al., 2017). Here we used the gait space classification methods outlined in (Wilshin et al., 2018) to test the similarity between proposed and measured gait patterns. Inspired by earlier work that used unsupervised learning to create ethograms from kinematic features (Hsu and Yttri, 2021), we combined these methods with clustering to identify patterns formed by the data in gait distance space, with the goal of identifying locomotory behaviors pre- and post-autotomy and regeneration that could either correspond to proposed gaits or new behaviors not previously reported (e.g., new gaits or novel compensatory mechanisms).

Finally, we investigated whether spiders use adaptive control and error-based learning in their locomotor response to limb loss, in which case we expected to see a time-dependent recovery of running performance after each autotomy (e.g., a monotonic return to pre-autotomy speed and a more stable gait). By contrast, if their neural control systems are robust to such perturbations, we expected to see a rapid recovery of stable locomotion by near-immediate adoption of the more stable gait (Mongeau et al., 2024). Consequently, we tested whether specimens exhibited a faster return to pre-autotomy speeds and more rapidly adopted the modified tripod gait after the second autotomy than the first autotomy.

## Methods

### Animals

Our spiders were captive-bred Guatemalan tiger rump tarantulas (*Davus pentaloris*, family *Theraphosidae)*. We purchased 45 spiderlings (2nd-3rd instar, 3-5 months old) from a private breeder (Mikolaz Sarnecki, Prospiders, Warsaw, Poland), ensuring that they had never lost limbs prior to our study. This species was chosen because they are fast runners, allowing us to assess learning while executing a high-performance behavior. The juveniles also molt and regenerate any autotomized limbs within 1-2 months. All specimens were allowed to acclimate in their new setting for at least 30 days before being used in experiments. Spiders were housed in an animal care room (25.6-27.8 deg C, 30% humidity, 12:12 light:dark schedule) and fed dubia (*Blaptica dubia*) and orange head (*Eublaberus posticus*) cockroaches. Only healthy specimens that were uninjured apart from the intended autotomies were used for experiments. The body length, *BL*, defined as the distance between the anterior end of the chelicerae to the caudal end of the abdomen (Fig. 2B), was measured on a calibrated photograph for each day on which experiments were performed (median *BL* 16.0 mm, range [7.4, 22.8] mm).

**Fig. 2.**
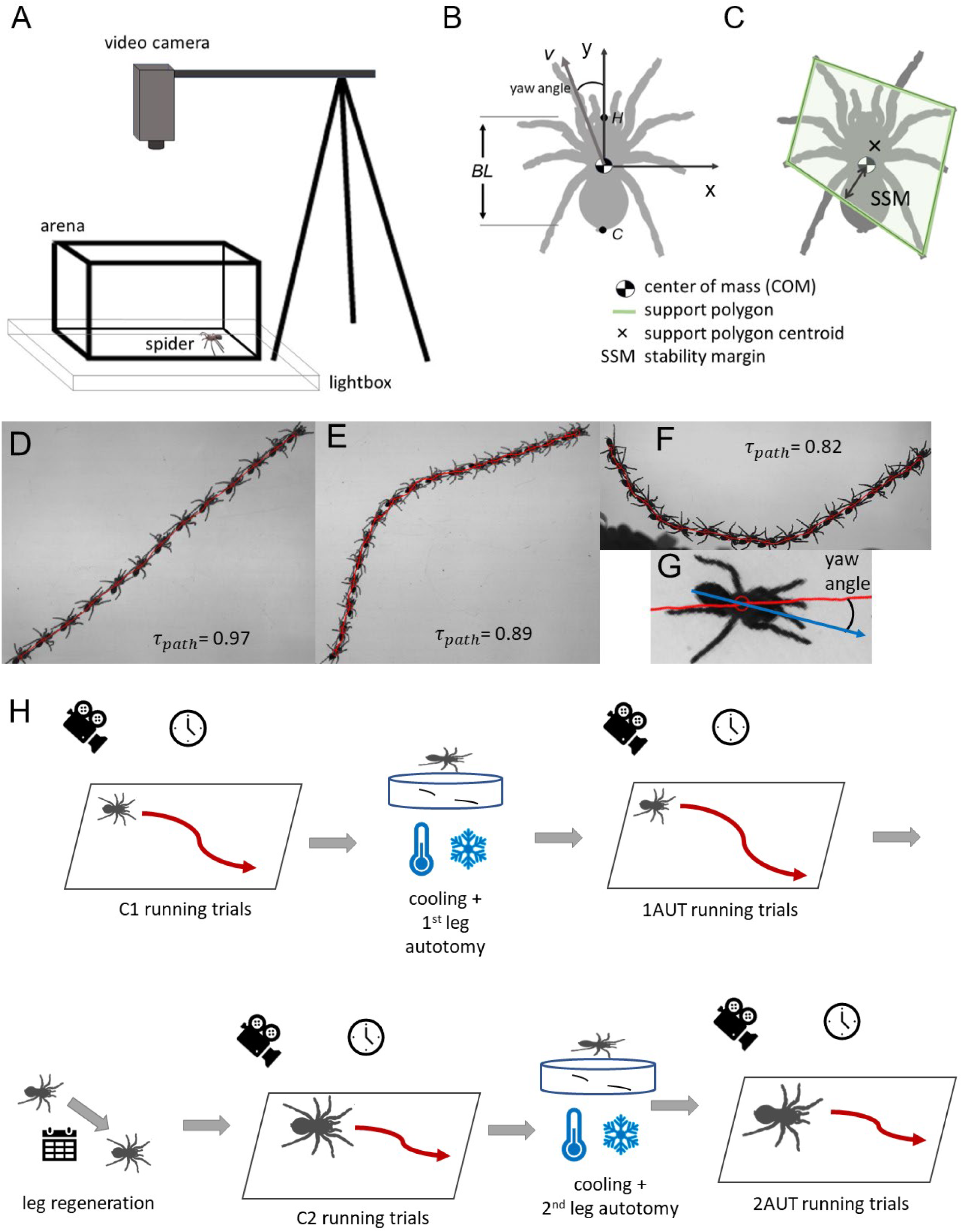
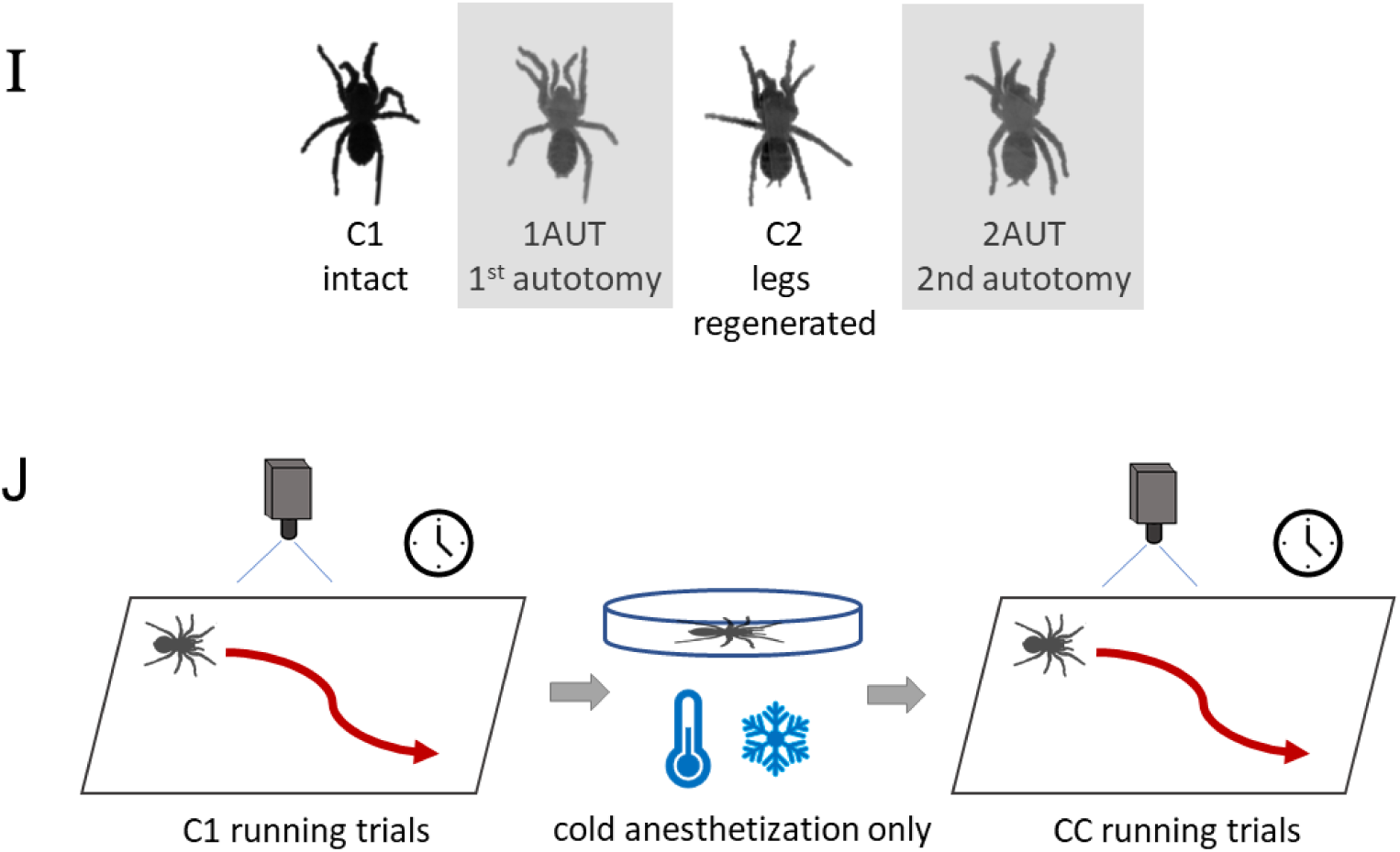
A) Schematic diagram of arena and filming geometry during running trials. B) Schematic illustration of the spider’s body length, *BL*, center of mass (COM), yaw angle between the cranial-caudal axis and center of mass velocity (*v*) and *xy* axes used for tracking motion in the comoving body frame. C) Illustration the definition of the support polygon (green polygon) and static stability margin (SSM). D-F) Superimposed images of running spiders filmed in dorsal view with the tracked body center of mass trajectory shown as a red line for different values of tortuosity, τ. G) Sample image of the yaw angle deviating from the velocity direction. H) Flow diagram showing steps in the protocol for the filming and experimental treatments. I) Still images from video of the same specimen after each treatment in H; gray shaded regions indicated autotomized specimens. J) Flow diagram of the cooling-only controls CC.

### Data analysis and statistics

All data analysis and statistics were performed using MATLAB v2023b (Mathworks, Natick MA USA) using the machine vision, curve fitting, statistics and machine learning toolboxes, unless stated otherwise; MATLAB functions are indicated below in italics. Differences between multiple means were computed using Kruskal-Wallis tests with post-hoc correction using Dunnett’s test (*kruskalwallis* and *multcompare* in MATLAB). Wilcoxon rank sum testing (*ranksum*) was used to compare paired measures. Linear regression was performed using *lmfit*. The reported p-values for linear regression fits were corrected using the Bonferroni-Holm post-hoc test for multiple comparisons (Holm, 1979) using the MATLAB implementation by (Groppe, 2024). All testing used a corrected significance level of 0.05. Results are reported as means [95% CI]; SD was used as a measure of variability, unless the Kolmogorov-Smirnov test for normality was inconsistent with a normal distribution, in which case we report the median and MAD (median absolute deviation). Bootstrap calculations were performed using the MATLAB function *bootci* using 1000 bootstrap samples. Gait calculations involving circular statistics were performed using the MATLAB toolbox CircSTAT (Berens, 2009).

### Experimental procedure

Videos were filmed using a single camera (Photron FASTCAM model SA-3, San Diego CA USA; settings: 8-bit monochrome, resolution 216 x 928 pixel, 500 frame s^−1^, timestep Δt = 2 ms/frame, shutter 1/1000 s, working distance 80 cm). The spiders were filmed in dorsal view during running in an indoor filming arena that consisted of a clear, smooth acrylic box (19 x 27 cm) at ambient temperature (24.0 ± 0.5 deg C) between 9:00 to 18:00. (Fig. 2A) The arena was placed on a lightbox that provided backlit illumination to eliminate shadows. The *xy* coordinates in the image plane corresponded to motions projected onto the horizontal plane in the laboratory. (Fig 2C). The video field of view showed spiders moving for a mean of 6 strides per trial (range [2, 14]), where the stride period was defined as the time required for one full cycle of foot motion during locomotion. Spiders were stimulated to move by approaching with a stick to provoke an escape response and allowed to run until they either stopped, reached the edge of the arena, or otherwise left the camera’s field of view. Here a “trial” refers to a single recorded bout of locomotion. After each trial, we retrieved the spider and quickly initiated a new trial until it ceased to move. While videos were being saved, the spider was placed under a small petri dish to limit its movement. After each set of trials was completed, the substrate was cleaned to remove possible silk and scent deposits that might serve as sensory cues for other spiders (Persons et al., 2001).

Five individual spiders (N = 5) were subjected to two successive autotomy treatments that involved the loss of two legs (hindleg L4 and foreleg R1). (Fig. 1C, D, 2H, I) Following (Wilshin et al., 2018), these legs were chosen because they belong to the same tetrapod in the alternating tetrapod gait, so their loss was expected to induce maximum gait disruption. In addition, forelegs and hindlegs have been reported to be the most frequently autotomized legs in several field studies of spiders (Johnson and Jakob, 1999; Steffenson et al., 2014; Wrinn and Uetz, 2007).

Fig. 2H illustrates the various experimental treatments. (Dataset 1) Control trials were performed on specimens with all legs intact prior to undergoing autotomy (C1). Spiders were then induced to self-autotomize in response to two of their legs being restrained, a natural process that occurs frequently in the field. In the protocol for the first autotomy treatment, each spider first was cold-anesthetized on ice at −1 deg C in a covered Petri dish containing a small piece of cardstock onto which the spider was placed. The dish was removed from ice once the spider was unresponsive to stimulation by touching (12-25 min), and the tarsus of each leg to be autotomized was adhered to the card using cyanoacrylate glue. Upon recovery, the spider quickly shed the two restrained legs. Each spider was placed in the arena within 46 s of autotomy and filmed repeatedly until it stopped running; this treatment corresponded to the closest match to the experiments on autotomized wolf spiders in (Wilshin et al., 2018). To allow a determination of the time course of any changes in locomotion, we recorded the time elapsed from the moment of limb loss to the start of each video recording. After the first round of experiments on the day of autotomy (1AUT0), the specimen was replaced in its habitat, allowed to move, eat, and drink freely overnight, and then recorded running on the following day using the same protocol as the first day to ascertain running performance after being allowed to recover for an additional 19-24 hr (1AUT1).

After both days of filming were complete, the specimens were allowed to regenerate their lost legs. We examined each regenerated leg to make sure it was fully developed (i.e., the same length, thickness, and morphology as the intact leg on the opposite side) and allowed the exoskeleton to harden for 1-2 weeks after molting before performing further experiments on the specimen. For four of the five specimens, we were able to record trials with the spider running using the regenerated legs (C2). Finally, for all five specimens we autotomized the same two legs as in the first autotomy experiments and recorded additional locomotion trials using the same protocol as after the first autotomy on the day of autotomy (2AUT0) and 20-32 hr after autotomy (2AUT1).

We also performed experiments to test the effect of cold-anesthetization on spider locomotion. This involved first filming a regular set of control trials (C1, N = 6) at ambient temperature, using only specimens that had not undergone autotomy. These specimens were then subjected to the same cooling treatment used before autotomy, but without inducing leg loss, then filmed while running using the standard protocol used for autotomized specimens (cooled controls, CC; N = 6). (Fig. 2J) Because these cooled controls necessarily were performed on specimens before the other treatments, shorter times were required to anesthetize smaller, younger specimens for cooled controls than for larger, older specimens prior to autotomy.

### Video analysis and kinematics calculations

The anatomical features tracked on video are shown on Fig 1B and 2B. Tracked landmarks comprised the spider’s body binary center of mass (COM) on video (which agreed with the location of the COM measured mechanically in (Biancardi et al., 2011; Silva-Pereyra et al., 2019)), the cranial and caudal ends (which also provided a measure of the body length, *BL*), and the distal tips of all tarsi. The *xy* coordinates of each landmark were measured first by custom automated tracking code written in MATLAB (see details in Appendix S1) followed by semiautomated manual tracking in Direct Linear Transformation data viewer (DLTdv) (Hedrick, 2008) to check all tracked data, correct any mistracked points and add any missing tracks. The spatial calibration (3.67 ± 0.02 pixel/mm) was measured from images of a ruler recorded for each recording session. The tracking uncertainty was ± 0.8 mm for *xy* coordinates and ± 2 deg for the orientation angle of the cranial-caudal axis.

For the majority of the C1 controls, CC cooling controls, and autotomy treatments, a total of five trials per specimen were analyzed. (Note that here we use the terms video and trial interchangeably.) The following exceptions were due to specimens becoming reluctant to run: 1) only 4 trials were recorded for one specimen’s CC treatment; 2) trials were recorded for only four of the five specimens for the C2 control after leg regeneration (five trials for two specimens, four trials for the other two).

For the 1AUT0 and 2AUT0 treatments and for the CC cooled controls, we analyzed videos recorded at times ranging from <1 to 44 min after the specimen experienced leg loss (for the autotomy experiments) or was removed from the cooling treatment (for cooled controls). We analyzed videos recorded at similar times after the start of each experiment for C1 and C2 controls, and for 1AUT1 and 2AUT1 (recorded the day after autotomy).

The tracked coordinates were used to compute several kinematic measures to compare running performance before and after leg loss, leg regeneration and cooling. The *xy* velocities of the COM, ***v***, and of each tarsus, ***v****_i_*, were computed from the tracked coordinates using a running quadratic fit (D’Errico, 2024) to the data over a time window of 50 ms for the COM and 40 ms for the tarsal coordinates; these times were chosen after initial testing found that smoothing the original data over these times using a running quadratic fit (*smooth* with the ‘rloess’ option) results in changes no greater than the tracking uncertainty. Previous research has found that spider running speed varies as body mass, M, as *v_*COM*_* ∝ *M*^0.353 ±0.08^ (Boehm et al., 2021) and that body mass scales with body length as *M* ∝ *BL*^2.70 ±0.04^ (Penell et al., 2018). We therefore expected that running speed should scale approximately linearly with body length: *v_*COM*_* ∝ *BL*^0.95±0.22^. In agreement with this, linear regression found that the measured mean running speed varied linearly with body length during control trials (R-squared = 0.80; v = 21.0 [19.0, 23l0] ms^−1^ × BL; slopes agreed at the 95% CI for C1 and C2 data). Consequently, relative speed (BL/s) was reported to compensate for variability in body length, as in previous studies of spider locomotion (e.g., Spagna and Peattie, 2012; Silva-Pereyra *et al*., 2019). To look for evidence of steering failure, we measured path tortuosity using the straightness index *τ*_*path*_ = *d*;⁄*D* ≤ 1, where *d* = displacement (i.e., distance between the initial and final COM positions) and *D* = total distance traveled along the trajectory (McLean and Skowron Volponi, 2018). In addition, to look for heading error, in which the body axis is misaligned with the direction of motion (Meshkani et al., 2023), we measured the yaw angle, defined as the angle between the instantaneous direction of COM velocity and the cranial-caudal axis (Wilshin et al., 2018). (Fig. 2B, G)

The motion of each intact tarsus was tracked in order to quantify gait kinematics. For each leg involved in locomotion, the gait cycle was defined as the time from when a specific tarsus first makes contact with the ground in a gait cycle until the instant before the same tarsus touches down in the next gait cycle. The stride period, *T_stride_*, and frequency, *f_stride_* = 1/ *T_stride_*, for each tarsus’s motion were defined as the mean spacing of successive peaks in the tarsus’s velocity along the COM motion. To show tarsal motion during locomotion, footfall diagrams were created for each time interval during which the COM speed was approximately constant. (Fig. 5) Each tarsus’s motion during a stride were divided into a “stance” state when the tarsus was fixed relative to the ground and a swing state when the tarsus moved relative to the ground. The stance state for each tarsus was estimated from the fraction of each gait cycle in which it moved caudally with respect to the COM; the swing state was then defined as the remainder of the gait cycle. The duty factor was estimated as the percent of frames in each gait cycle in which the tarsus in question was in stance. The stride frequency and stride length were computed for each tarsus, then averaged over all tarsi.

### Stability measures

We also computed several measures of static stability previously used in analyzing quasi-static locomotion (Ting et al., 1994). To do so, we first computed for each video frame the estimated number of tarsi in the stance state, *N_support_*, and the support polygon, defined as the convex hull of the *xy* coordinates of all tarsi in stance. (Fig. 2C) For static stability, the support polygon must at minimum be a tripod (*N_support_* ≥ 3) (Ting et al., 1994). The static stability margin, *SSM*, was determined from the minimum distance from the support polygon’s edges to the projection of the COM onto the ground. For an organism at rest with respect to the ground that exerts only downward ground reaction forces, the system is statically stable when *SSM* > 0 and in metastable equilibrium when *SSM* = 0. If the centroid of the support polygon is at the same location as the COM, then *SSM* is at its maximum value, the ideal static stability margin *ISSM*, and the system is said to be ideally statically stable; more generally *SSM* ≤ *ISSM*. For *SSM* < 0, an organism at rest is statically unstable. In the analysis of this study’s data, we interpreted these stability measures cautiously because running tarantulas do not necessarily move quasi-statically. Even if SSM < 0, they can still locomote stably by relying on dynamic stability mechanisms (Koditschek et al., 2004) or by grasping and adhering to surfaces (Silva et al., 2021).

To optimize stability, autotomized spiders could adopt an altered range of leg motions adjusted to cover the functional space of both the intact and missing tarsi. To explore this possibility, we measured the average angle between adjacent legs. If the legs responded to leg loss to remain uniformly spaced in angle around the COM, then the angle between adjacent legs should increase from 45 deg = 360 deg / 8 for intact specimens to 60 deg = 360 deg / 6 after autotomy, consistent with the measured values. Second, the distribution of each tarsus’s range of lateral and fore-aft motions provided a measure of whether neighboring tarsi filled in the functional space left vacant by an autotomized tarsus. To determine whether any tarsi (e.g., those adjacent to each missing leg (L3 next to autotomized L4 and R2 next to autotomized R1) increased their range of motion to cover the functional space covered by two legs in the intact specimen, we plotted the position of each tarsus in a body-fixed frame with the +y-axis oriented along the COM velocity. (Fig. 2B) In this frame, we measured the area swept out by the range of motion of each tarsus, after omitting outliers (i.e., values > 3 MAD from the median), for intact and autotomized specimens. Third, because a wider stance provides a greater stabilizing torque for the same ground reaction force, we computed the fore-aft and lateral distances between each tarsus and the COM (leg extensions) for pre- and post-autotomy.

### Gait analysis

Gait patterns were computed and then analyzed using unsupervised machine learning and compared to proposed model gaits: the alternating tetrapod (ALT) for intact spiders with 8 legs, and the ablated tetrapod (ABT) and modified tripod (MT) gaits for autotomized spiders with 6 legs.

First, gait patterns were quantified using the methods developed in (Revzen and Guckenheimer, 2008) to compute the phase of oscillation of each leg during locomotion. This approach is a generalization of relationships for a simple harmonic oscillator moving periodically along the y-axis with constant amplitude *A* and frequency, *ω*. For the solution in which position at time *t*, *y* = *Acosωt*, then velocity, 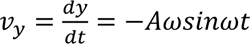, and phase, 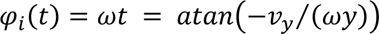. More generally, during locomotion the position, y_*i*_, and velocity, v_*yi*_ of *i* th tarsus are oscillatory signals with noisy amplitude and frequency, and the effective oscillation phase can be estimated as:

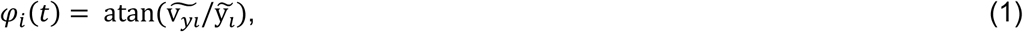

where position, 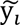, and velocity, 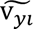, have been standardized to zero mean and unit variance (Revzen et al., 2013). (Fig. 3A) The difference in oscillation phase between each pair of intact adjacent legs, Δ𝜑_*i*_, is computed as:

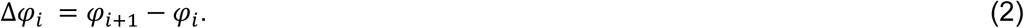

in which the intact legs are numbered *i* = 1, 2, …, *N_legs_* starting from the foremost left leg and running counterclockwise in dorsal view to the foremost right leg, as shown in Fig. 3A. Because 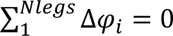, we define each gait using (*N_legs_* – 1) independent values of phase differences: Δ𝜑*_*gait*_* = (Δ𝜑_1_, Δ𝜑_2_, …, Δ𝜑*_*Nlegs*_*_− 1_). For example, for an intact spider, the alternating tetrapod gait is defined by Δ𝜑_𝐴𝐿𝑇_ = (Δ𝜑_1_, Δ𝜑_2_, …, Δ𝜑_7_), with all Δ𝜑_*i*_ = 0.5 in a 7-dimensional gait space. Similarly, for autotomized spiders with 6 legs, the modified tripod has Δ𝜑_*M*𝑇_ = (Δ𝜑_1_, Δ𝜑_2_, …, Δ𝜑_5_) with all Δ𝜑_*i*_ = 0.5 in a 5-dimensional gait space. The ablated tetrapod gait differs from the modified tripod in that the third phase difference is 0 instead of 0.5: Δ𝜑_𝐴𝐵𝑇_ = (Δ𝜑_1_, Δ𝜑_2_, …, Δ𝜑_5_) = (0.5, 0.5, 0, 0.5, 0.5).

**Fig. 3.**
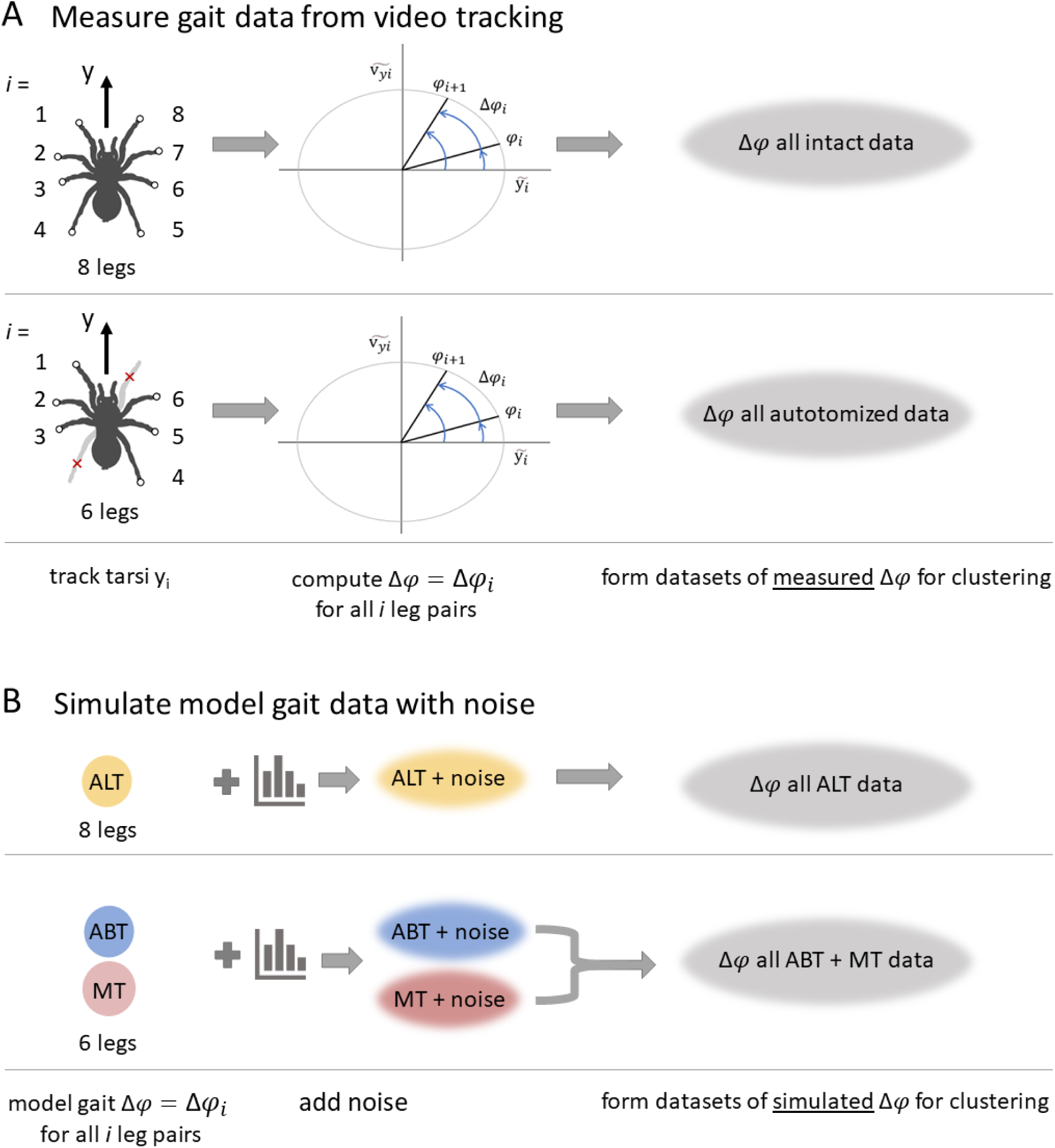
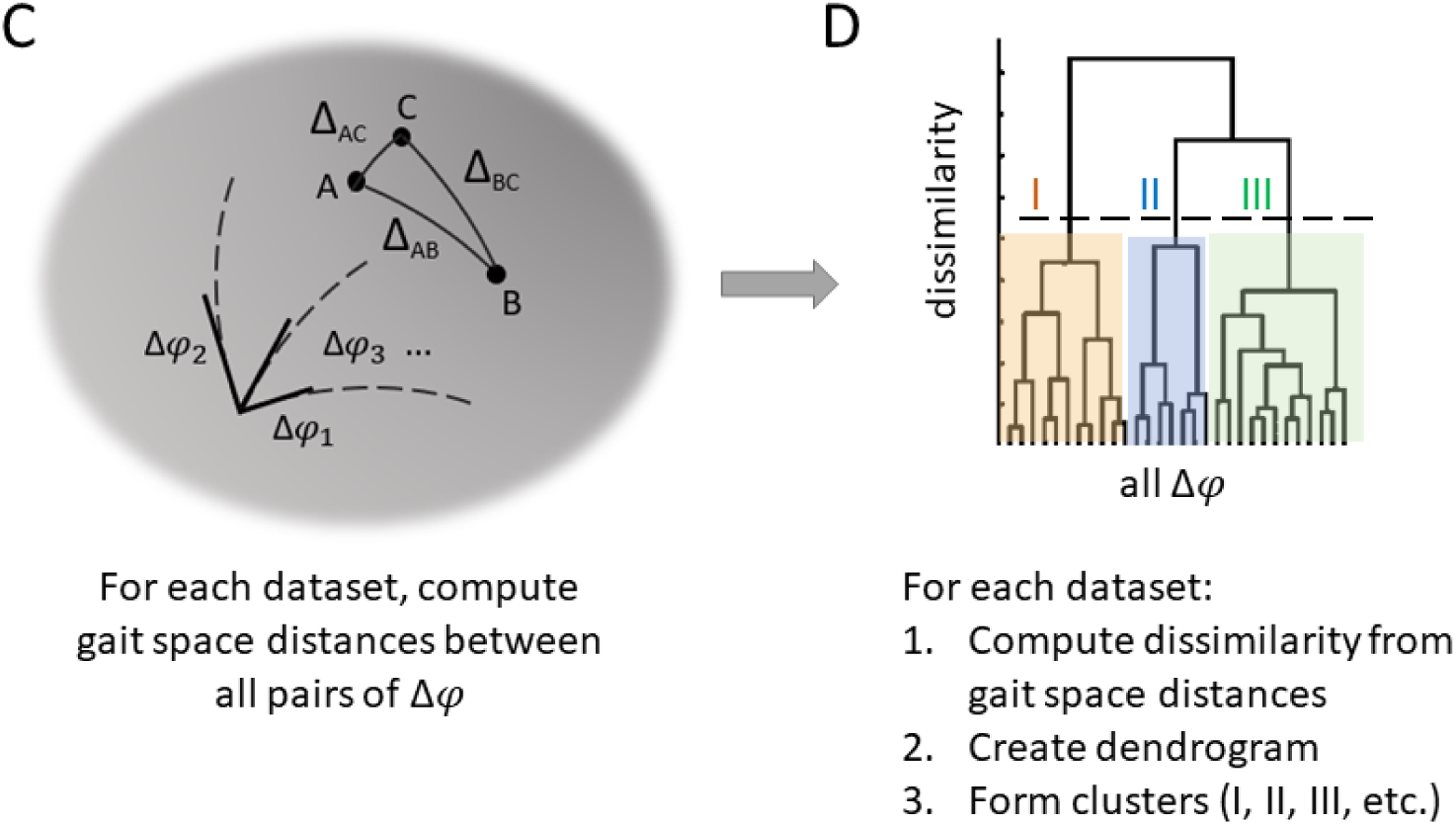
Flow diagram illustrating the pipeline for calculating and clustering measured and simulated gait phase differences for autotomized treatments. A) Illustration of the calculation of the *i* th leg’s phase, 𝜑_*i*_, (Eq. 1), the phase difference, Δ𝜑_*i*_, between the *i* th pair of adjacent legs (Eq. 2) from standardized values of the position, 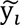, and velocity, 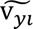, of each tarsus, and the formation of datasets for intact and separately for autotomy treatment data. B) Simulated gaits were created with added noise for these models: ALT = alternating tetrapod, ABT = ablated tetrapod, and MT = modified tripod. The ABT and MT data then were combined to form a simulated autotomy dataset for clustering. Four datasets of phase differences were created for clustering as described in A and B. C) These datasets of phase differences were then each used to compute the distances, Δ_AB_, between each pair of data points in the (N_legs_-1)-dimensional non-Euclidean gait space. D) Finally, the distances were used to compute a dendrogram based on the dissimilarity between pairs of points. Clusters were assigned based on branches in the dendrogram with the greatest dissimilarities.

To quantify measured gait patterns, first tracked values of the *i* th tarsus’s position and velocity along the fore-aft direction (y_*i*_ and v_*yi*_) were used in Eqn 1 and 2 to compute the (*N_legs_* −1) phase differences, Δ𝜑_*i*_, for each frame. All values of phase and phase differences were computed in units of oscillation cycles. Next, these data were analyzed using their positions in a metric gait space with gait coordinates Δ𝜑*_*gait*_* = (Δ𝜑_1_, Δ𝜑_2_, …, Δ𝜑*_*Nlegs*_*_−1_); for full details, see (Wilshin et al., 2017; Wilshin et al., 2018). Because each of the Δ𝜑_*i*_ is periodic with range [0, 1] cycle, each gait corresponds to a point on an (*N_legs_* – 1) dimensional hypertorus. (Fig. 3D) The distance between two points in this gait space, *d_AB_*, corresponding to two different gaits A (Δ𝜑_𝐴_) and B (Δ𝜑_𝐵_) depends on the minimum distance, 𝛿𝛿_𝐴𝐵*i*_, along each dimension Δ𝜑_*i*_ for all *i* = 1, 2, …, (*N_legs_* – 1) leg pairs:

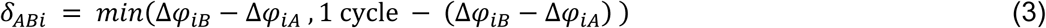

This is because each value of 𝛿_𝐴𝐵*i*_ can be thought of as the minimum distance between two points on a circle with unit circumference. As explained in (Wilshin et al., 2018), the total distance *d_AB_* between points A and B in gait space then can be computed from the set of 𝛿_𝐴𝐵*i*_ for all *i* leg pairs by using a metric tensor, *g_ij_*, (Table S1) that accounts for the mean phase of all legs during a gait cycle and ensures each *i* th leg pair’s phase difference is equally weighted. The gait distance, *d_AB_*, in cycles is then defined by:

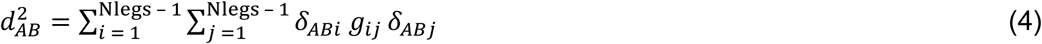

The maximum distance possible between two points in gait space, *d*;_𝑚𝑎𝑥_ = 𝑚𝑎𝑥 *d*;_𝐴𝐵_, occurs when 𝛿_𝐴𝐵*i*_ = 0.5 cycle for all *i* (Table S1). We used this fact to compute a normalized gait distance Δ_AB_ that has range [0,1]:

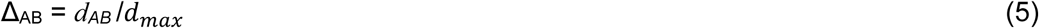

For example, the modified tripod and ablated tetrapod gaits are separated by a distance Δ_ABT-MT_ = 0.29 of the maximum possible distance. To compare a dataset of measured phase differences (e.g., all Δ𝜑_*i* 𝑓𝑟𝑎𝑚𝑒_ for each frame of each trial for a given treatment) to the Δ𝜑_*i gait*_ for a model gait, we used Eq. 5 along with bootstrapping to measure the mean and 95% CI of the gait space distances between all measured and model phase differences. This allowed us to compare, for example, gait patterns measured for intact spiders with the ideal alternating tetrapod gait, and those for autotomized spiders with both the ideal ablated tetrapod and modified tripod gaits.

We also extended the gait space methods by building on ideas for ethogram creation using unsupervised learning (Hsu and Yttri, 2021). In this approach, datasets of measured leg phase differences, Δ𝜑_*i*_, for all intact treatments and separately for all autotomized treatments were assigned to groups (hereafter, clusters) using hierarchical agglomerative clustering and Ward’s linkage (*linkage*, *dendrogram* and *cluster* in MATLAB) with the gait space distances precomputed using Eq. 5 as the similarity measure. (Fig. 3C, D) This allowed us to group together more or less similar values of Δ𝜑_*i*_ without reference to predefined gaits. Different clusters of measured data were compared to each other and to hypothesized model gaits using bootstrapped values of mean and 95%CI gait space distance.

To validate using unsupervised learning to group noisy measured data, we also performed clustering on synthetic gait data with simulated noise. We started by creating a perturbed gait dataset for each of the ideal model gaits. (Fig. 3B) For each model gait, we generated 1000 random samples of Δ𝜑_*i*_ with added noise using the von Mises distribution (*circ_vmrnd* (Berens, 2009)) with the parameter μ_i_ that characterizes the average equal to the Δ𝜑_*i*_ values for the model gait and the mean dispersion parameter κ determined from fitting the measured phase difference data for the intact and separately for the autotomized specimens to a von Mises distribution to the. The simulated perturbed ALT data then were compared to the results for the intact measured data. To create a simulated dataset for comparison with the measured autotomy data, we merged the perturbed ablated tetrapod and modified tripod samples into a single dataset. (Fig. 3B) These two simulated sets of perturbed data were then analyzed using the gait space distance and clustering methods described above. (Fig. 3C, D)

The results of clustering the measured and simulated data were used to estimate the effect of noise on both phase differences and gait space distances. This approach allowed us to explore whether the leg motions during locomotion corresponded to a series of successive motions corresponding to a hypothesized model gait, a different, but still regular, gait pattern, or a noisy series of leg motions not corresponding to a fixed gait pattern per se. We also used the clustering results to compare statistics for the speed, the static stability margin and the number of feet in stance for the different gait patterns identified.

## Results

All kinematic and gait analyses described below were performed on a total of 43562 frames, equivalent to 96-173 strides per treatment (Dataset 1). The spiders consistently ran without stumbling or falling, as shown in the typical videos for C1, C2 and 1AUT0 treatments in Movie 1. We first consider whether various locomotion measures depended on the time after treatment. Body yaw angle, tortuosity, stride frequency, duty factor and static stability margins did not depend significantly on time in any trials, and running speed had a significant correlation with time in only 1 of 24 trials (statistics in Dataset 1). We therefore pooled together the data for each trial across times before performing further analysis.

### Running performance

We next tested the effect on running performance of the two successive two-leg autotomies and subsequent regeneration. Fig. 4A-C shows the results of comparing the mean speed, tortuosity and yaw for the control C1 to other treatments. (For full statistics, see Table S2, S3). Tarantula running speed did not differ significantly between controls conducted before and after leg regeneration (C1 vs C2). There also was no reduction in speed associated with autotomy: while speed and stride frequency were reduced relative to the control on the day of autotomy (but not the day after), this effect was not significantly different from the decrease due to cold anesthetization alone (CC) (Fig. 4A, D). There were no significant differences in stride length among treatments (Fig. S1D).

**Fig. 4.**
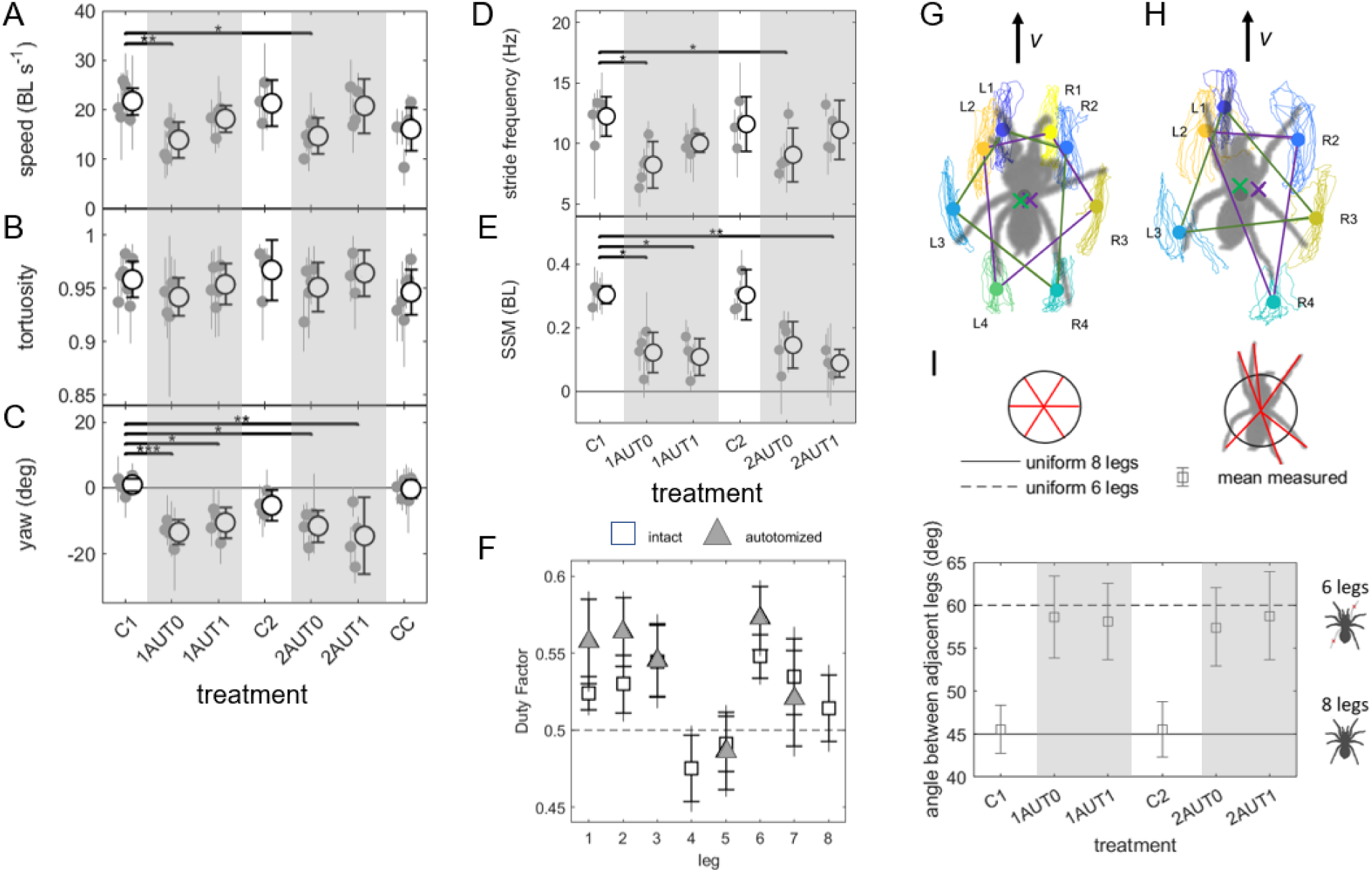
Kinematic and posture measurements among treatments. A-E) show summary statistics for each variable for each treatment as grand means or medians and 95% CI (black open circles and error bars); gray filled markers and error bars show means or medians and 95% CI for each specimen. (For each treatment, N = 5 specimens, n = 5 trials/specimen.) Gray shaded regions correspond to autotomy treatments. Treatments that are significantly different from the controls C1 as indicated by Kruskal-Wallis testing with correction are indicated by lines and * or ** (P ≤ 0.05 or 0.01, respectively). F) The duty factor for each leg for the intact and autotomy treatments (grand means, error bars: 95%CI (with caps) and SD (no caps)). G, H) Examples of typical tarsal range of motion during locomotion. Colored lines and markers indicate measured tarsus range-of-motion trajectories and motion centroids for G) an intact specimen with 8 legs (C1 control) and H) the same specimen with 6 legs the day after the first autotomy treatment (1AUT1). In these figures, the distal end of each tarsus is plotted in a body-fixed reference frame in which the body COM (gray circle) is at the origin and the velocity, **v**, is oriented vertically. The mean positions of each tarsus (colored circles) in the corresponding tetrapods (G) and tripods (H) are shown as purple and green × markers. I) Comparison of the angle between adjacent legs for each treatment expected if legs are uniformly distributed in angle about the center of mass (solid circles) with the measured grand mean and 95%CI (open circles and error bars).

We separately tested whether the spiders had improved running performance for treatment 2AUT0 compared to 1AUT0, consistent with their learning from the first autotomy, using Wilcoxon ranked sum testing between paired treatments of speed, tortuosity, and yaw grouped by specimen. None of these measures differed between the first and second autotomy.

The path tortuosity, *τ*_*path*_, did not differ significantly among any treatments. (Fig. 4B) The tortuosity varied over a narrow range close to the maximum value of 1 for a straight line (135 values in the range [0.896, 0.993], one outlier at 0.82) (Fig. 2D-F).

Fig. 4E shows that the static stability margin, SSM, decreased significantly in all but one autotomy treatment. For intact and autotomized treatments, the duty factor exceeded 0.5 (i.e., the tarsus was in contact with the ground for over half the stride cycle on average) for all but the hindmost legs (i.e., legs *i* = 4 and 5; see Fig. 4F). This is because the hindlegs frequently drag (see videos for the intact and regrown cases in Movie 1); during dragging, tarsi that rest on the ground move, thereby lowering their duty factor, even though they are able to support weight.

We next consider evidence for postural changes post-autotomy. During pre-autotomy controls C1 and CC, the spiders oriented their cranial-caudal axis along the velocity direction (zero yaw) on average. By contrast, after autotomy, the specimens ran with their bodies rotated 11-15 deg relative to the forward velocity. (Fig. 4C) See Fig. 4G, H for a typical example of how the motions and mean orientations of the different legs compared before and after autotomy. To test whether this change in yaw was accompanied by differences in leg use, we compared measured angles between adjacent legs with the values expected if the legs were uniformly spaced in angle. (Fig. 4I) For every control and treatment considered, the theoretical and measured angles between legs agreed, consistent with autotomized spiders increasing the mean angle between adjacent legs until they were spread uniformly about the COM. Fig. S1A-C shows the results of tests for whether the tarsi moved through a greater range or extended more widely laterally and fore-aft after autotomy. The only significant difference in their range of motion area was a 1.3× increase after the first autotomy. (Fig. S1A) The tarsi did not extend significantly farther in the fore-aft and lateral directions after autotomy and leg regeneration. (Fig. S1B, C)

### Gait analysis

Fig. 5 shows the level of agreement between footfall diagrams for the idealized gaits proposed for the intact and autotomized spiders and for measured data for trials with the minimum and median gait distances to each idealized gait. Good visual agreement is apparent between the measured and idealized diagrams for all but that for the trial with the median distance to the ablated tetrapod (Fig. 5H). To explore these gait patterns further, we next consider the results of clustering data from the gait space analysis for the simulated and measured phase differences.

**Fig. 5.**
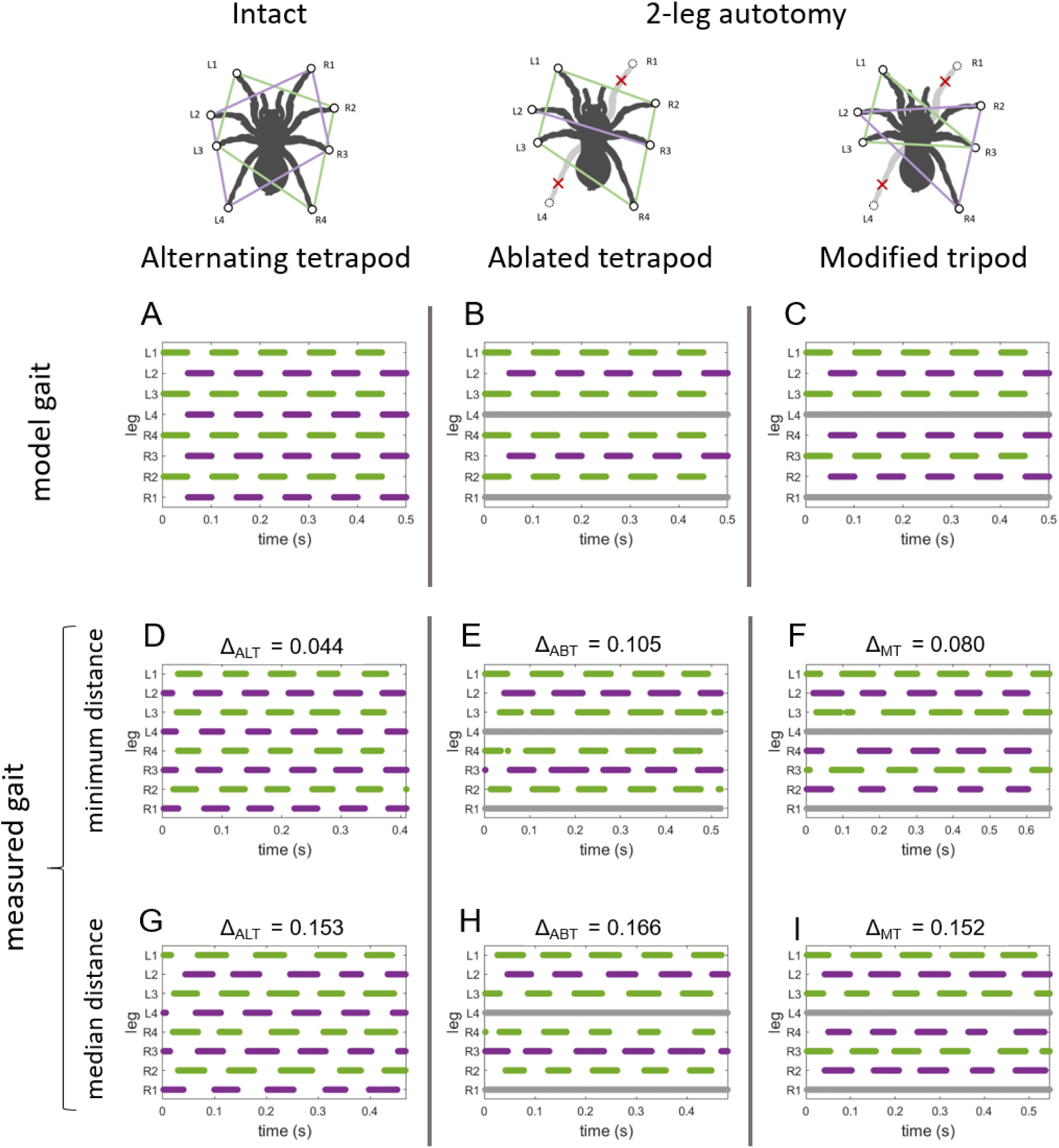
Gait footfall diagrams for idealized models and measured data. The corresponding pods (sets of coordinated legs) are shown schematically at the top as polygons in colors matching their hypothesized motion; gray lines indicate missing data for autotomized legs. Simulated model gaits are shown in A-C. Examples computed from empirical data are shown for 5 strides/trial for trials with minimum gait distances (D-F) and for trials with the grand median distance from the closest model gait (G-I). The gait space distance to the closest model gait is given above each diagram. The corresponding treatments were: C1 for D, G; 1AUT0 for E, F; 1AUT1 for I; 2AUT1 for H.

First, we found that the synthetic data created by adding noise to the ablated tetrapod and modified tripod were correctly grouped with the original model gait by clustering for 95% of samples; assignment by closest gait distance was correct in 92% of samples. Given this validation of the accuracy of clustering applied to simulated gait data, we next applied this method to measured data.

We created two datasets for the cluster analysis by pooling the gait space data for: 1) all intact control trials (C1, C2); and 2) all of the autotomy trials (1AUT0, 1AUT1, 2AUT0, 2AUT1), because Kruskal-Wallis testing showed no significant differences among treatments (Dataset 1). Fig. 6A and B show the two dendrograms created by applying hierarchical clustering to each of these empirical phase difference datasets. These dendrograms display the height (dissimilarity) derived from the gait distance computed for each pair of leaves shown at the bottommost end of each branch; here, a leaf is a measured set of Δ𝜑_*i*_, for all *i* leg pairs. Each branch corresponds to a cluster of gait patterns with varying level of similarity. To find the clusters with the greatest dissimilarity, we start at the top of each dendrogram and find the branch points at the greatest height. For the dendrogram formed by the intact data, this corresponds to the branch point between cluster ALT3 and a second branch; continuing downward, this second branch splits into clusters ALT1 and ALT2. (Fig. 6A) (The names for these and all other gait clusters were based on their proximity between the clusters’ centroids and the relevant model gaits; e.g., for intact data, clusters ALT2, ALT1 and ALT3 have centroids that are 11.5%, 14.6% and 20.4% of the maximum possible distance from the alternating tetrapod. (Fig. 6C)) Similarly, the dendrogram for the autotomy data first branches into cluster ABT1 and a second branch, which further splits into the branches labeled MT1 and MT2 (Fig. 6B) (Statistics in Dataset 1). Based on the similarity of the gait space and kinematic analysis of clusters ALT1 and ALT2 and clusters MT1 and MT2 for the intact and autotomy data, respectively, described below, we did not further subdivide the data. For the autotomy data, the MT1 and MT2 clusters are closer to the simulated modified tripod (MT) data, and the ABT1 cluster is closer to the ablated tetrapod (ABT) simulation. (Fig. 6D)

**Fig. 6.**
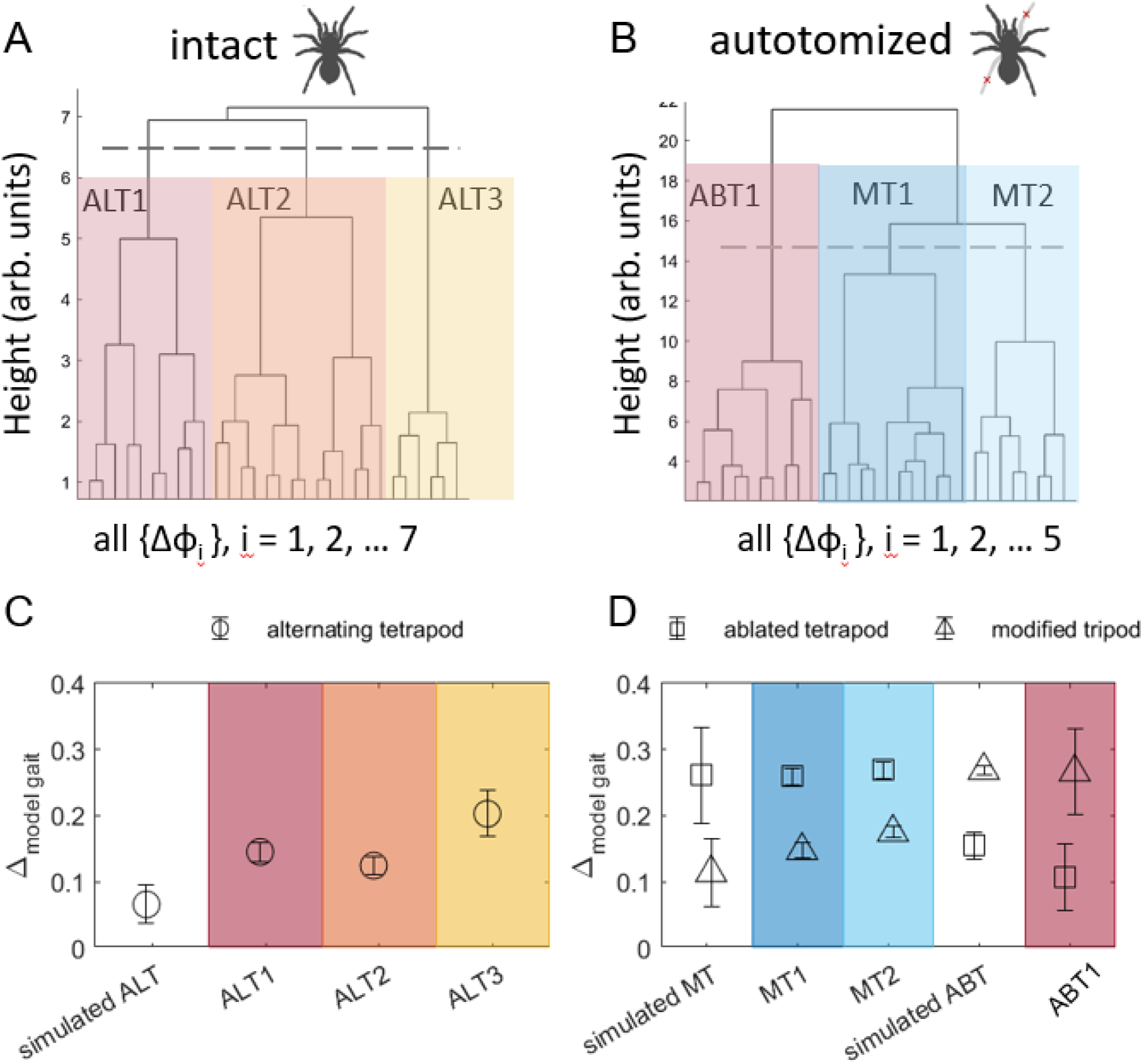
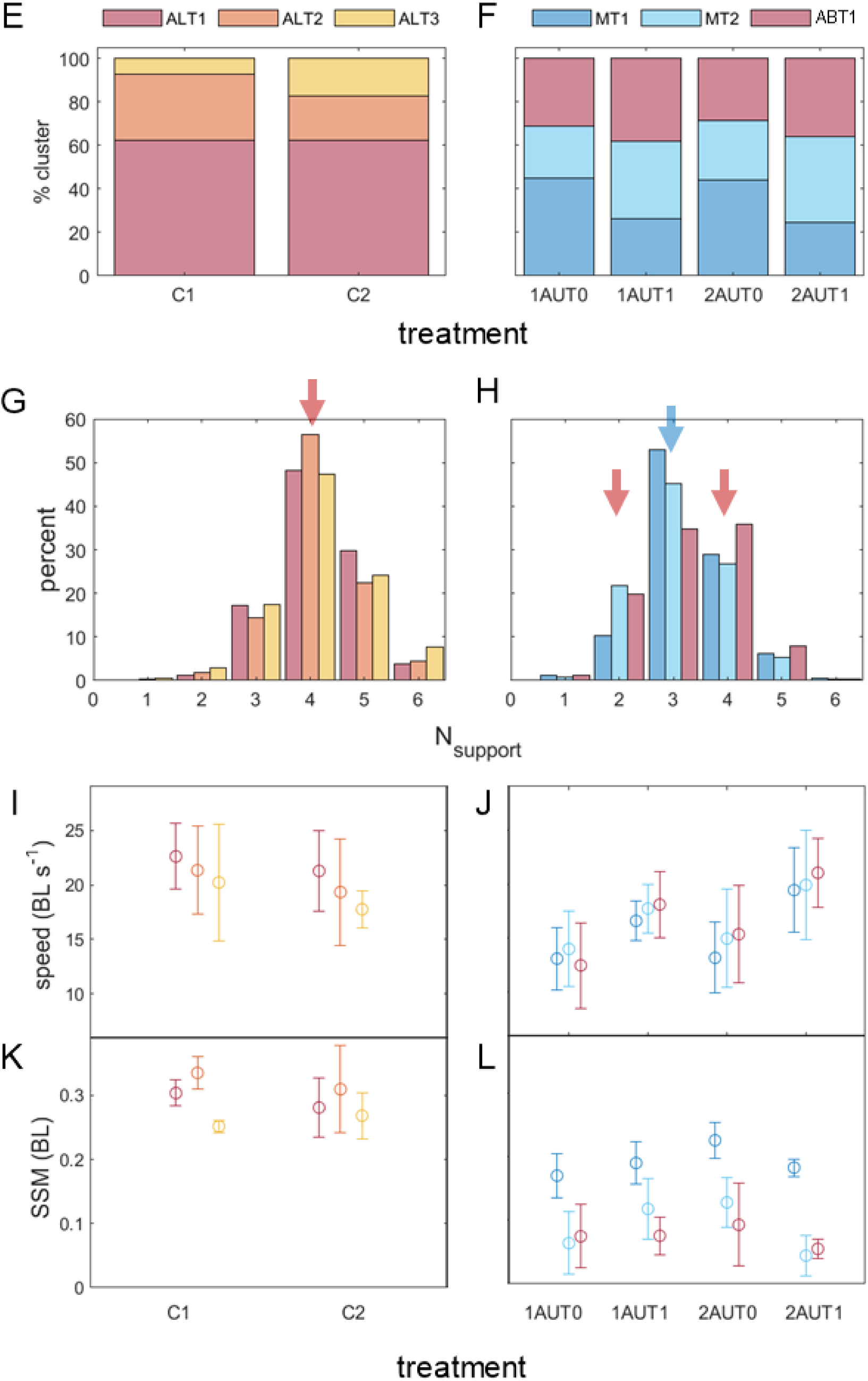
Gait space analysis. Dendrograms for A) all intact (C1 and C2 control) data and B) all autotomized treatment data (1AUT0, 1AUT1, 2AUT0, 2AUT1). Leaves in the dendrogram correspond to measured phase space differences, while heights correspond to dissimilarity between the data computed using gait space distances. Clusters are labeled by the most similar model gait: ALT1, 2, 3 for alternating tetrapod, ABT1 for ablated tetrapod, and MT1, 2 for modified tripod. Comparison of gait space distances from the ideal model gaits for C) intact and D) autotomy simulations and measured gait clusters (grand means, SD error bars). (For each treatment, N = 5 specimens, n = 5 trials/specimen.) E, F) Fractions of the data grouped into the clusters shown in A and B by treatment averaged over specimens. G, H) Histograms indicate the percent of time for which each number of tarsi was in stance (*N_support_*) by cluster for all intact and all autotomized treatments. Colored arrows indicate the expected values for the relevant model gaits: magenta at *N_support_* = 4 for the alternating tetrapod in G, and *N_support_* = 2 and 4 for the ablated tetrapod in H, blue for the modified tripod in H. Comparison of the I, J) speed (grand mean, SD error bars), K, L) static stability margin, SSM (grand median, MAD error bars). (Marker and bar colors in I-L agree with those used in C-H.)

The distributions of clusters are shown in Fig. 6E, F; chi-squared testing indicated significant differences in the distributions among all treatments. (Dataset 1) The majority (62%) of the intact data corresponded to ALT1, with the distribution between the other clusters changing from 7.5% of C1 in ALT3 to 17.5% of C2. (Fig. 6E) The majority of gait data for autotomized treatments (Fig. 6F) was predominantly associated with clusters similar to the modified tripod (MT1 and MT2). To test whether learning resulted in the spiders adopting gaits similar to the modified tripod to a greater extent at later times, we compared the fraction of data clustered as either MT1 or MT2 (MT clusters) for autotomy treatments at different times. The fraction of data in MT clusters decreased by 7.3% from the first to second day of each autotomy treatment (i.e., from 1AUT0 to 1AUT1 and from 2AUT0 to 2AUT1), but slightly increased (2.2%) between the first and second autotomy treatments on either day (i.e., from 1AUT0 to 2AUT0 and from 1AUT1 to 2AUT1).

Fig. 6I, J show that the speed for each cluster was similar among treatments. In particular, the speeds for each cluster in the autotomy data were similar for the 1AUT0 and 2AUT0 treatments, indicating no effect of learning due to prior experience with autotomy.

Comparing the static stability margin, SSM, among clusters revealed that the MT2 and ABT1 clusters had a lower SSM than MT1. (Fig. 6 K, L) Distributions for treatments for intact specimens have a dominant peak at N_support_ = 4, as expected for the alternating tetrapod. (Fig. 6M) For autotomy treatments (Fig. 6N), the MT1 and MT2 cluster data had a single peak at N_support_ = 3, the value expected for the modified tripod gait. However, the ABT1 cluster had its most prominent values at N_support_ = 3 and 4; while 4 was consistent with one intact tetrapod (4), the peak at 3 disagreed with the value of 2 expected for the bipod formed by the ablated set of two tarsi. Thus, even though the ABT1 cluster was closer in gait space to the ablated tetrapod model gait, the spiders in question adjusted the timing of their leg motions so they only had two or fewer tarsi on the ground < 27% of the time. This shortened stance phase for the bipod was referred to as “limping” when used by wolf spiders in (Wilshin et al., 2018). For comparison, the corresponding value for the intact data was < 2%. In general, none of the data for any treatment agreed with an aerial phase (i.e., intervals during which N_support_ = 0 and hence no tarsi are in contact with the ground,).

Fig. 7 shows the distribution of measured phase differences, Δ𝜑_*i*_, for each leg pair for each cluster. In the intact data, the hindmost legs (Δ𝜑_3_, Δ𝜑_4_, Δ𝜑_5_) moved out of phase by 0.5 cycle, in agreement with the alternating tetrapod. However, for each ipsilateral pair of forelegs (i.e., (Δ𝜑_1_, Δ𝜑_2_, Δ𝜑_6_, Δ𝜑_7_), the phase of the more cranial of each pair, 𝜑_*i*_, lagged that of the other leg. For the autotomized data, the phase differences between remaining leg pairs (Δ𝜑_1_, Δ𝜑_2_, Δ𝜑_4_) were similar to the values in the intact data. The phase differences affected by an autotomized leg (Δ𝜑_3_, Δ𝜑_5_) were in close agreement with values for the relevant model gaits. In general, there were similar spreads for all leg pairs for all treatments and clusters, with the exception of a larger spread in Δ𝜑_3_ between the legs adjoining the autotomized hindleg for the ABT1 cluster.

**Fig. 7.**
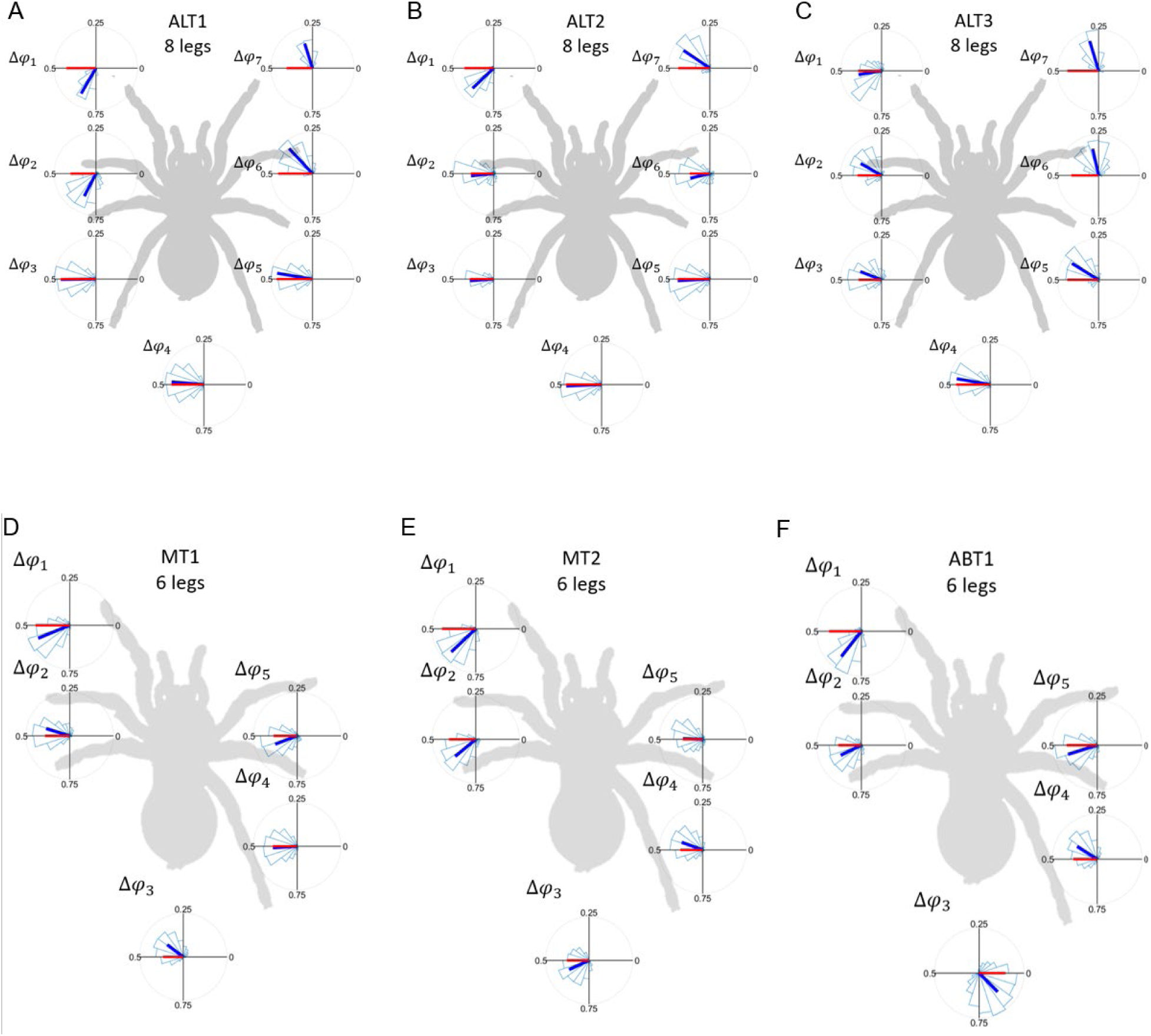
Polar plots of measured leg pair phase difference histograms. Circular statistics are shown for the leg pair phase differences, for A-C) the three clusters (ALT1, ALT2, ALT3) for all the intact data and D-F) the three clusters (MT1, MT2 and ABT1) for all the autotomy treatments. Polar plots of histograms (light blue open bars) of the distributions of the phase differences, Δ𝜑_*i*_, between pairs of adjacent legs for all trials for all autotomized trials grouped into the three clusters indicated in A, B). In each plot, the direction of the circular mean vector is shown by a blue solid line; a red solid line corresponds to the direction of the most similar model gait. The length of both lines indicates the amount of dispersion of the measured Δ𝜑_*i*_, with shorter lines corresponding to greater dispersion. If the direction and length of the blue and red lines agree, this indicates good agreement between the measured and model gaits (statistics in Dataset 1).

To illustrate how these results corresponded to dynamic leg motions, we created videos that showed the spider in a frame of reference with the COM fixed and the vertical aligned with the fore-aft direction, with the tarsi in stance in each frame indicated by a support polygon. (Movie 1). For each frame, we also plotted a label indicating the cluster assignment and colored the support polygon according to the most similar set of comoving tarsi for the corresponding model gait (Fig. 1B-C). These videos illustrate variable gait patterns that did not conform to two stable sets of legs alternately placed in stance. We also observed that for the autotomy treatments, frames in which only a statically unstable bipod or monopod were in stance often were followed quickly by more cranially-oriented tarsi being planted to form support polygon.

## Discussion

Juvenile spiders that had never experienced autotomy were found to achieve a stable gait immediately after 25% leg loss and to resume their pre-autotomy speed within one day. In addition, two-leg autotomy following cold anesthetization did not result in a speed significantly lower than due to cold anesthetization alone in the hour after treatment; this lack of an additional effect due to leg loss was surprising because the autotomy treatment was designed to maximally impact their normal gai pattern. Furthermore, the spiders did not increase their path tortuosity in response to leg loss, unlike harvestmen and water-walking insects (Escalante and O’Brien, 2024; O’Neil et al., 2024). Overall, this indicates these spiders achieved a faster recovery of running performance than the two days found for autotomized harvestmen (Escalante et al., 2020). A comparison of data for the first and second autotomy treatments did not support a role for learning in this recovery: there was no significant difference between kinematic measures and only a small (2.2%) change in gait use (i.e., between gaits similar to the more stable modified tripod vs less stable ablated tetrapod). These results agree with a picture in which spiders respond to the loss of legs by rapidly adjusting their locomotion to achieve a new stable gait (robust control), rather than using error-based learning (adaptive control) (Mongeau et al., 2024).

While autotomy did not affect their legs’ extension or range of motion, the spiders did modify their posture so as to move with nonzero body yaw and more widely-spread legs. Spiders with regenerated legs also ran at the same speed as before autotomy, as found for purple shore crabs (Prestholdt et al., 2022).

The gait clusters found by applying unsupervised learning to the interleg leg phase difference data approximated the ideal model gaits proposed for intact and autotomized spiders (Wilshin et al., 2018; Wilson, 1967). Furthermore, similar variability was found in the phase differences between most pairs of adjacent legs. However, these clusters also exhibited additional structure in the measured gait patterns that differed from these models. For example, among other discrepancies, all of the identified clusters had several adjacent leg pair differences that disagreed with the predicted 0.5 or 0 cycle offsets of the model gaits, indicating that the legs did not move in two synchronized groups as predicted by the model gaits. These variations resulted in more than one cluster resembling a model gait in some cases. For example, the data for intact spiders could be subdivided into clusters with differing levels of agreement with the alternating tetrapod. In these gait clusters, the ipsilateral and contralateral pairs of adjacent hindlegs swung out of phase as predicted by the alternating tetrapod gait, but the foreleg pairs were closer to being in phase than expected. In another example, the autotomized data had two clusters that resembled variants of the modified tripod (MT1 and MT2) with different timing of leg motions, as well as different static stability margins.

This complexity relates to a known challenge in locomotor gait analysis: the fact that leg motions often are too irregular and complicated to be described fully by static patterns of phase relationships between legs. Examples of irregular gaits in octopods are shown and discussed in, e.g., (Biancardi et al., 2011; Silva-Pereyra et al., 2019; Spagna and Peattie, 2012; Weihmann, 2013; Wilson, 1967). Earlier studies had defined diverse gaits used for locomotion by intact spiders based on mean phase differences between leg pairs (Biancardi and Silva-Pereyra, 2020; Weihmann, 2020).

Here, the use of clustering to group leg motions enabled us to meet this challenge by identifying patterns in the gait data corresponding to differences in both regular coordination and variability in the timing of leg motions. This gait analysis was interpreted in comparison with the kinematic results and videos annotated to show both the clustering results and leg motions. These findings indicate that these spiders adapted to leg loss by adopting gaits in which a specific pattern of leg motion is not rigidly followed. The phase difference distributions (Fig. 7) indicate that for most clusters the measured interleg coordination patterns between pairs of the hindmost legs were the most consistent, and those involving the forelegs the least consistent, with values expected for the corresponding model gaits. This may relate to their different roles, in that forelegs play a role in pulling, sensing, and deceleration, while hindlegs primarily propel and stabilize (Pullar and Paulin, 2018). We also observed the spiders dragging one or more hindlegs, which contributed to the finding that their duty factor < 0.5 in some frames and may serve to enhance stability (Spagna et al., 2011; Weihmann et al., 2015). In addition, the distribution of feet in stance at any time did not follow expectations from model gaits. The observed deviations from the model’s 0.5 cycle phase difference resulted in the foremost legs having duty factors > 0.5 and more legs in stance at any given time than predicted for alternately-moving pods of feet. Notably, this resulted in the cluster most similar to the ablated tetrapod gait (ABT1) most frequently having 3 and 4 feet in stance, avoiding the model’s prediction of a 50:50 frequency of 4 for the tetrapod and 2 for the statically unstable bipod. The ABT1 and MT2 clusters both had a higher instance of two or fewer tarsi on the ground than MT1, resulting in a lower static stability margin. However, the similar speed found for all three clusters for autotomy treatments can be explained if these spiders are dynamically stable as defined in (Hof et al., 2005). For example, humans walking with crutches can achieve this type of dynamic stability in the swing-through gait, allowing them to undergo stable inverted pendulum motion about a pivot point formed alternately by the two crutch tips and the “good leg” (Rasouli and Reed, 2020). The annotated videos for autotomized specimens show examples in which short sequences of a statically unstable bipod is followed by a stable support polygon (Movie 1); in these cases, the bipod serves as a temporary pivot point about which the spider rotates quickly to plant additional feet on the ground to maintain stable motion, the behavior referred to as “limping” in (Wilshin et al., 2017).

More generally, gait space clustering can identify patterns in empirical data without reference to predefined gaits, allowing the matching of measured gaits with known patterns as well as the discovery of new locomotor behaviors. This meets a pressing need in the study of locomotion by multi-legged animals, for which automated tracking now facilitates the collection of extremely large datasets of leg motions that are not readily interpretable using standard methods. To facilitate applying these methods in other contexts, we provide code for analyzing the gait space distances and performing clustering on phase data for four-, six- and eight-legged locomotion that can be modified to work with varying degrees of leg loss. This approach could be used to determine how locomotion varies during different behaviors (e.g., moving at different speeds or orientations, carrying loads, pursuing prey, or displaying) or in response to other challenges, such as differences in terrain or changes in environmental conditions. Because gait data alone cannot identify behaviors such as leg dragging, vertical body motions, and movements that enable dynamic stability, ideally it should be analyzed in combination with kinematic data and annotated video. In future work, these methods can be modified to work with three-dimensional data for leg and body motions facilitated by deep learning methods for automatically tracking motion on video (O’Neil et al., 2024). This should enable, for example, relating changes in gait to the mechanical work performed during locomotion (Silva-Pereyra et al., 2019) and how animals redistribute loads among their remaining legs in response to leg loss (Meshkani et al., 2023).

The comparative biomechanics of limb autotomy is also an important source of inspiration for legged robot design. Past studies of how robots can achieve stable gait coordination with one or more damaged or missing legs have used arthropod-inspired control algorithms and sensors that allow reprogramming gait phase relationships as well as body and leg posture (Görner and Hirzinger, 2010; Mostafa et al., 2010; Schilling et al., 2007; Shih et al., 2012; Spenneberg et al., 2004; Zhang et al., 2024), and leg compliance (Görner and Hirzinger, 2010). Using clustering to provide a deeper understanding of actual gait patterns during animal locomotion promises further insights relevant for robotics.

In conclusion, we found that intact and autotomized spiders both ran at similar speeds using variable gait patterns that only approximated model gaits in which legs move in rigidly-synchronized sets. This study also found that that body orientation interacts with leg use, with an increased body yaw facilitating the redistribution of leg angles required to adapt the limb kinematics to compensate for leg loss. Together these adaptations appear to render spiders robust to the perturbations caused by leg loss, and thereby likely more responsive to similar challenges encountered during routine locomotion, such as uneven terrain, missed footing, and using some legs for nonlocomotory purposes (e.g., load-bearing or sensing). These results are relevant for understanding how arthropods cope with limb loss and for designing fault-tolerant locomotion in robotics.

## Data availability

All data and code required to reproduce the results and figures presented here are provided in the Supplementary Information as Dataset 1.

## Acknowledgements

We wish to thank Simon Wilshin and Luis Contreras-Orendain for helpful conversations.

## Funding

NSF CAREER grant (IOS-1453106) to S.T.H; Temple Univ. Science Scholar Program Stipend to B.L.Q.

## Notes

https://figshare.com/articles/dataset/Dataset_S1_for_Unsupervised_learning_reveals_rapid_gait_adaption_after_leg_loss_and_regrowth_in_spiders/28229174

### Competing Interest Statement

The authors have declared no competing interest.

## References

Apontes, P. and Brown, C. A. (2005). Between-sex Variation in Running Speed and a Potential Cost of Leg Autotomy in the Wolf Spider Pirata sedentarius. The American Midland Naturalist 154, 115–125.

Barnes, W. J. P. (1975). Leg co-ordination during walking in the crab,Uca pugnax. J. Comp. Physiol. 96, 237–256.

Berens, P. (2009). CircStat: A MATLAB Toolbox for Circular Statistics. Journal of Statistical Software 31, 1–21.

Biancardi, C. M. and Silva-Pereyra, V. (2020). Biomechanics of Locomotion in Tarantulas. In New World Tarantulas: Taxonomy, Biogeography and Evolutionary Biology of Theraphosidae (ed. Pérez-Miles, F.), pp. 365–388. Cham: Springer International Publishing.

Biancardi, C. M., Fabrica, C. G., Polero, P., Loss, J. F. and Minetti, A. E. (2011). Biomechanics of octopedal locomotion: kinematic and kinetic analysis of the spider Grammostola mollicoma. Journal of Experimental Biology 214, 3433–3442.

Boehm, C., Schultz, J. and Clemente, C. (2021). Understanding the limits to the hydraulic leg mechanism: the effects of speed and size on limb kinematics in vagrant arachnids. J Comp Physiol A 207, 105–116.

Bowerman, R. F. (1975a). The control of walking in the scorpion. I. Leg movements during normal walking.

Bowerman, R. F. (1975b). The control of walking in the scorpion. II. Coordination modification as a consequence of appendage ablation.

Brown, C. A. and Formanowicz, D. R. (2012). The effect of leg autotomy on terrestrial and aquatic locomotion in the wolf spider Pardosa valens (Araneae: Lycosidae). The Journal of Arachnology 40, 234–239.

Brown, C. A., Amaya, C. C. and Formanowicz, D. R. (2018). The frequency of leg autotomy and its influence on survival in natural populations of the wolf spider *Pardosa valens*. Can. J. Zool. 96, 973–979.

Brueseke, M. A., Rypstra, A. L., Walker, S. E. and Persons, M. H. (2001). Leg Autotomy in the Wolf Spider Pardosa milvina: A Common Phenomenon with Few Apparent Costs. amid 146, 153–160.

D’Errico, J. (2024). Movingslope. MATLAB Central File Exchange.

Emberts, Z., Escalante, I. and Bateman, P. W. (2019). The ecology and evolution of autotomy. Biological Reviews 94, 1881–1896.

Escalante, I. and O’Brien, S. L. (2024). Robustness to Leg Loss in Opiliones: A Review and Framework Considerations for Future Research. Integrative and Comparative Biology icae051.

Escalante, I., Badger, M. A. and Elias, D. O. (2020). Rapid recovery of locomotor performance after leg loss in harvestmen. Sci Rep 10, 13747.

Fleming, P. A., Muller, D. and Bateman, P. W. (2007). Leave it all behind: a taxonomic perspective of autotomy in invertebrates. Biological Reviews 82, 481–510.

Foelix, R. F. (2011). The biology of spiders. Third edition. Oxford University Press.

Gerald, G. W., Thompson, M. M., Levine, T. D. and Wrinn, K. M. (2017). Interactive effects of leg autotomy and incline on locomotor performance and kinematics of the cellar spider, Pholcus manueli. Ecology and Evolution 7, 6729–6735.

Görner, M. and Hirzinger, G. (2010). Analysis and evaluation of the stability of a biologically inspired, Leg loss tolerant gait for six- and eight-legged walking robots. In 2010 IEEE International Conference on Robotics and Automation, pp. 4728–4735. Anchorage, AK: IEEE.

Groppe, D. (2024). Bonferroni-Holm Correction for Multiple Comparisons. MATLAB Central File Exchange.

Han, Q. (2024). Interleg coordination in free-walking bug Erthesina fullo (Hemiptera: Pentatomidae). Insect Science.

Hedrick, T. L. (2008). Software techniques for two- and three-dimensional kinematic measurements of biological and biomimetic systems. Bioinspir. Biomim. 3, 034001.

Herreid, C. F., II and Full, R. J. (1986). Locomotion of Hermit Crabs (Coenobita Compressus) on Beach and Treadmill. Journal of Experimental Biology 120, 283–296.

Hof, A. L., Gazendam, M. G. J. and Sinke, W. E. (2005). The condition for dynamic stability. Journal of Biomechanics 38, 1–8.

Holm, S. (1979). A Simple Sequentially Rejective Multiple Test Procedure. Scandinavian Journal of Statistics 6, 65–70.

Hsu, A. I. and Yttri, E. A. (2021). B-SOiD, an open-source unsupervised algorithm for identification and fast prediction of behaviors. Nat Commun 12, 5188.

Hughes, G. M. (1957). The Co-Ordination of Insect Movements: II. The Effect of Limb Amputation and the Cutting of Commissures in the Cockroach (Blatta Orientalis). Journal of Experimental Biology 34, 306–333.

Jayaram, K. (2015). Robustness of Biological and Bio-inspired Exoskeletons.

Johnson, S. A. and Jakob, E. M. (1999). Leg autotomy in a spider has minimal costs in competitive ability and development. Animal Behaviour 57, 957–965.

Koditschek, D. E., Full, R. J. and Buehler, M. (2004). Mechanical aspects of legged locomotion control. Arthropod Structure & Development 33, 251–272.

Land, M. F. (1972). Stepping Movements Made by Jumping Spiders During Turns Mediated by the Lateral Eyes. Journal of Experimental Biology 57, 15–40.

Lutzy, R. M. and Morse, D. H. (2008). Effects of leg loss on male crab spiders *Misumena vatia*. Animal Behaviour 76, 1519–1527.

McLean, D. J. and Skowron Volponi, M. A. (2018). trajr: An R package for characterisation of animal trajectories. Ethology 124, 440–448.

Meshkani, J., Rajabi, H., Kovalev, A. and Gorb, S. N. (2023). Locomotory Behavior of Water Striders with Amputated Legs. Biomimetics 8, 524.

Mongeau, J.-M., Yang, Y., Escalante, I., Cowan, N. and Jayaram, K. (2024). Moving in an Uncertain World: Robust and Adaptive Control of Locomotion from Organisms to Machine Intelligence. Integrative and Comparative Biology icae121.

Mostafa, K., Tsai, C.-S. and Her, I. (2010). Alternative Gaits for Multiped Robots with Leg Failures to Retain Maneuverability. International Journal of Advanced Robotic Systems 7, 34.

O’Neil, J. N., Yung, K. L., Difini, G., Rohilla, P. and Bhamla, M. S. (2024). Limb loss and specialized leg dynamics in tiny water-walking insects. 2024.04.02.587762.

Penell, A., Raub, F. and Höfer, H. (2018). Estimating biomass from body size of European spiders based on regression models. arac 46, 413–419.

Persons, M. H., Walker, S. E., Rypstra, A. L. and Marshall, S. D. (2001). Wolf spider predator avoidance tactics and survival in the presence of diet-associated predator cues (Araneae: Lycosidae). Animal Behaviour 61, 43–51.

Pfeiffenberger, J. A. and Hsieh, S. T. (2021). Autotomy-induced effects on the locomotor performance of the ghost crab Ocypode quadrata. Journal of Experimental Biology 224, jeb233536.

Prestholdt, T., White-Toney, T., Bates, K., Termulo, K., Reid, S., Kennedy, K., Turley, Z., Steed, C., Kain, R., Ortman, M., et al. (2022). Tradeoffs associated with autotomy and regeneration and their potential role in the evolution of regenerative abilities. Behavioral Ecology 33, 518–525.

Pullar, K. F. and Paulin, M. G. (2018). Markerless tracking suggests a tactile sensing role for forelegs of Dolomedes spiders during locomotion. 398479.

Rasouli, F. and Reed, K. B. (2020). Walking assistance using crutches: A state of the art review. Journal of Biomechanics 98, 109489.

Revzen, S. and Guckenheimer, J. M. (2008). Estimating the phase of synchronized oscillators. Phys. Rev. E 78, 051907.

Revzen, S., Burden, S. A., Moore, T. Y., Mongeau, J.-M. and Full, R. J. (2013). Instantaneous kinematic phase reflects neuromechanical response to lateral perturbations of running cockroaches. Biol Cybern 107, 179–200.

Saintsing, A. J. (2022). Robustness in Locomotion: The Energetic Cost of Perturbations.

Schilling, M., Cruse, H. and Arena, P. (2007). Hexapod Walking: an expansion to Walknet dealing with leg amputations and force oscillations. Biol Cybern 96, 323–340.

Shih, T.-S., Tsai, C.-S. and Her, I. (2012). Comparison of Alternative Gaits for Multiped Robots with Severed Legs. International Journal of Advanced Robotic Systems 9, 157.

Silva, V., Biancardi, C., Perafán, C., Ortíz, D., Fábrica, G. and Pérez-Miles, F. (2021). Giant steps: adhesion and locomotion in theraphosid tarantulas. J Comp Physiol A 207, 179–190.

Silva-Pereyra, V., Fábrica, C. G., Biancardi, C. M. and Pérez-Miles, F. (2019). Kinematics of male Eupalaestrus weijenberghi (Araneae, Theraphosidae) locomotion on different substrates and inclines. PeerJ 7, e7748.

Spagna, J. C. and Peattie, A. M. (2012). Terrestrial locomotion in arachnids. Journal of Insect Physiology 58, 599–606.

Spagna, J. C., Valdivia, E. A. and Mohan, V. (2011). Gait characteristics of two fast-running spider species (Hololena adnexa and Hololena curta), including an aerial phase (Araneae: Agelenidae). arac 39, 84–91.

Spenneberg, D., McCullough, K. and Kirchner, F. (2004). Stability of walking in a multilegged robot suffering leg loss. In IEEE International Conference on Robotics and Automation, 2004. Proceedings. ICRA ‘04. 2004, pp. 2159–2164 Vol.3.

Steffenson, M. M., Formanowicz, D. R. and Brown, C. A. (2014). Autotomy and its Effects on Wolf Spider Foraging Success. Ethology 120, 1128–1136.

Ting, L. H., Blickhan, R. and Full, R. J. (1994). Dynamic and static stability in hexapedal runners. Journal of Experimental Biology 197, 251–269.

Tross, J., Wolf, H., Stemme, T. and Pfeffer, S. E. (2022). Locomotion in the pseudoscorpion Chelifer cancroides: forward, backward and upside-down walking in an eight-legged arthropod. Journal of Experimental Biology 225, jeb243930.

Weihmann, T. (2013). Crawling at High Speeds: Steady Level Locomotion in the Spider Cupiennius salei—Global Kinematics and Implications for Centre of Mass Dynamics. PLOS ONE 8, e65788.

Weihmann, T. (2020). Survey of biomechanical aspects of arthropod terrestrialisation – Substrate bound legged locomotion. Arthropod Structure & Development 59, 100983.

Weihmann, T., Goetzke, H. H. and Günther, M. (2015). Requirements and limits of anatomy-based predictions of locomotion in terrestrial arthropods with emphasis on arachnids. Journal of Paleontology 89, 980–990.

Weihmann, T., Brun, P.-G. and Pycroft, E. (2017). Speed dependent phase shifts and gait changes in cockroaches running on substrates of different slipperiness. Front Zool 14, 54.

Wilshin, S., Reeve, M. A., Haynes, G. C., Revzen, S., Koditschek, D. E. and Spence, A. J. (2017). Longitudinal quasi-static stability predicts changes in dog gait on rough terrain. Journal of Experimental Biology 220, 1864–1874.

Wilshin, S., Shamble, P. S., Hovey, K. J., Harris, R., Spence, A. J. and Hsieh, S. T. (2018). Limping following limb loss increases locomotor stability. Journal of Experimental Biology 221, jeb174268.

Wilson, D. M. (1967). Stepping Patterns in Tarantula Spiders. Journal of Experimental Biology 47, 133–151.

Wrinn, K. M. and Uetz, G. W. (2007). Impacts of leg loss and regeneration on body condition, growth, and development time in the wolf spider *Schizocosa ocreata*. Can. J. Zool. 85, 823–831.

Zeng, M., Meng, C., Han, B., Li, Y., Yu, H., Fu, H. and Zhong, S. (2024). Gait Characteristics and Adaptation Strategies of Ants with Missing Legs. J Bionic Eng.

Zhang, Y., Zhang, H., Ma, W., Li, P. and Fang, D. (2024). Gait planning and adjustment for a hexapod walking robot with one leg failure. Australian Journal of Mechanical Engineering 0, 1–14.

