## Supplementary Information for "Unsupervised learning reveals rapid gait adaption after leg loss and regrowth in spiders"

#### Movie 1. Video sequences for controls, autotomy treatments and trials with regenerated legs

This video compilation features: 1) sample videos for the C1 intact controls, a 1AUTO treatment (day of first two-leg autotomy treatment) and C2 (after leg regeneration); 2) Simulated videos illustrate the various model gaits hypothesized for these various treatments; 3) example videos that show the actual leg motions used by spiders with 8 legs (intact) and after two-leg autotomy, annotated to show the gait clusters used and the legs in stance in each frame.

### Appendix S1. Image analysis and tracking

Automated tracking consisted of several stages. (Note: italics are used to indicate functions in MATLAB.) First the original video was used to track the body center-of-mass, approximated as the location of the pedicel (Biancardi and Silva-Pereyra, 2020). Raw videos were preprocessed by contrast enhancement (*imadjust*) and conversion to binary (*imbinarize)*. A new body-only image then was created using morphological closing (*imclose* with a disk-shaped structuring element, radius = 8 pixels) to remove the legs, pedipalps and spinnerets. The resulting body-only image was analyzed for each video frame using *regionprops* to find the binary centroid (COM) and the orientation angle, $\theta$, of the cranial-caudal axis angle with respect to the x-axis, calculated using the best fit ellipse to the body binary image. A comparison between the manually and automatically tracked COM coordinates gave a root-mean-square coordinate discrepancy of ± 3 pixel uncertainty that agreed with that estimated from manually tracking. The root-mean-square discrepancy between manually and automatically tracked orientation angles was ± 2 deg. The COM coordinates and orientation angles were used to compute velocity and acceleration vs time. The COM trajectory's contour length (total distance traveled) was computed using *arclength* (https://www.mathworks.com/matlabcentral/fileexchange/34871-arclength).

The time window used for computing time derivatives was determined by performing smoothing on the COM coordinate vs time data using *smooth* with the *rloess* option. This allowed us to determine that the original and smoothed coordinate data agreed to ≤ ± 3 pixels for smoothing windows ≤ 50 ms, comparable to larger arthropod response times (Hergenröder and Barth, 1983; Hesselberg and Vollrath, 2006; Nakata, 2009). We then used this time window for computing derivatives on unsmoothed coordinate data using *movingslope* (https://www.mathworks.com/matlabcentral/fileexchange/16997-movingslope), which performs a local quadratic fit over the specified time window.

A new video was created for appendage tracking by centering each frame's image on the COM, cropping to a view encompassing only the spider, and reorienting the image using *imrotate* such that the spider's cranial-caudal axis pointed along the y-axis. All analysis after this step therefore occurred in a coordinate frame comoving with the spider's body. Next, we performed image analysis to track the proximal and distal endpoints of each appendage (i.e., intact leg or pedipalp). This was accomplished by subtracting the best-fit ellipse to the body-only binary image, leaving only the appendages. Each appendages binary image was then "skeletonized" using *bwmorph* with the "thin" option, an operation that removes pixels from the boundary of objects so as to preserve their Euler number while reducing the object to lines. The appendage's skeleton was analyzed with *bwmorph* to find the xy coordinates of terminating vertices ("endpoints") and intersecting vertices ("branchpoints") of the resulting skeleton. Finally, we used kmeans clustering, Kalman filtering and Munkres' version of the Hungarian algorithm (MATLAB *assignDetectionsToTracks*) with the cost based on Euclidean distance to organize the tracked coordinate data into sets corresponding to each appendage's tarsus.

**References**

Biancardi, C.M. *et al.* (2011) 'Biomechanics of octopedal locomotion: kinematic and kinetic analysis of the spider Grammostola mollicoma', *Journal of Experimental Biology*, 214(20), pp. 3433–3442. Available at: https://doi.org/10.1242/jeb.057471.

Biancardi, C.M. and Silva-Pereyra, V. (2020) 'Biomechanics of Locomotion in Tarantulas', in F. Pérez-Miles (ed.) *New World Tarantulas: Taxonomy, Biogeography and Evolutionary Biology of Theraphosidae*. Cham: Springer International Publishing (Zoological Monographs), pp. 365–388. Available at: https://doi.org/10.1007/978-3-030-48644-0_13.

Bowerman, R.F. (1975a) ‘The control of walking in the scorpion. I. Leg movements during normal walking.’ Available at: https://www.cabidigitallibrary.org/doi/full/10.5555/19750528596 (Accessed: 23 March 2024).

Bowerman, R.F. (1975b) ‘The control of walking in the scorpion. II. Coordination modification as a consequence of appendage ablation.’ Available at: https://www.cabidigitallibrary.org/doi/full/10.5555/19750528597 (Accessed: 23 March 2024).

Hergenröder, R. and Barth, F.G. (1983) ‘The release of attack and escape behavior by vibratory stimuli in a wandering spider (Cupiennim salei keys.)’, *Journal of comparative physiology*, 152(3), pp. 347–359. Available at: https://doi.org/10.1007/BF00606240.

Hesselberg, T. and Vollrath, F. (2006) ‘Temperature affects both web spider response time and prey escape speed’, *BULLETIN-BRITISH ARACHNOLOGICAL SOCIETY*, 13(7), p. 275.

Nakata, K. (2009) ‘Attention focusing in a sit-and-wait forager: a spider controls its prey-detection ability in different web sectors by adjusting thread tension’, *Proceedings of the Royal Society B: Biological Sciences*, 277(1678), pp. 29–33. Available at: https://doi.org/10.1098/rspb.2009.1583.

Pullar, K.F. and Paulin, M.G. (2018) ‘Markerless tracking suggests a tactile sensing role for forelegs of Dolomedes spiders during locomotion’. bioRxiv, p. 398479. Available at: https://doi.org/10.1101/398479.

Silva-Pereyra, V. *et al.* (2019) ‘Kinematics of male Eupalaestrus weijenberghi (Araneae, Theraphosidae) locomotion on different substrates and inclines’, *PeerJ*, 7, p. e7748. Available at: https://doi.org/10.7717/peerj.7748.

Tross, J. *et al.* (2022) ‘Locomotion in the pseudoscorpion Chelifer cancroides: forward, backward and upside-down walking in an eight-legged arthropod’, *Journal of Experimental Biology*, 225(10), p. jeb243930. Available at: https://doi.org/10.1242/jeb.243930.

Weihmann, T. (2013) ‘Crawling at High Speeds: Steady Level Locomotion in the Spider Cupiennius salei—Global Kinematics and Implications for Centre of Mass Dynamics’, *PLOS ONE*, 8(6), p. e65788. Available at: https://doi.org/10.1371/journal.pone.0065788.

Wilshin, S. *et al.* (2018) ‘Limping following limb loss increases locomotor stability’, *Journal of Experimental Biology*, 221(18), p. jeb174268. Available at: https://doi.org/10.1242/jeb.174268.

Wilson, D.M. (1967) ‘Stepping Patterns in Tarantula Spiders’, *Journal of Experimental Biology*, 47(1), pp. 133–151. Available at: https://doi.org/10.1242/jeb.47.1.133.

Dataset 1. Code and datasets required to reproduce the results and figures

https://figshare.com/articles/dataset/Dataset_S1_for_Unsupervised_learning_reveals_rapid_gait_adaption_after_leg_loss_and_regrowth_in_spiders/28229174

Figure S1. Comparison of kinematic and posture measurements among treatments.

Black open circles and error bars indicate the grand mean and 95% CI for each treatment, while gray filled markers and error bars are means and 95% CI for each specimen. Treatments that are significantly different from the controls C1 as indicated by Kruskal-Wallis testing are indicated by lines and * or ** at the top for P ≤ 0.05 or 0.01, respectively; similarly, the results of Wilcoxon rank sum tests on paired 1AUT0 and 2AUT0 data are indicated by lines and asterisks at the bottom.

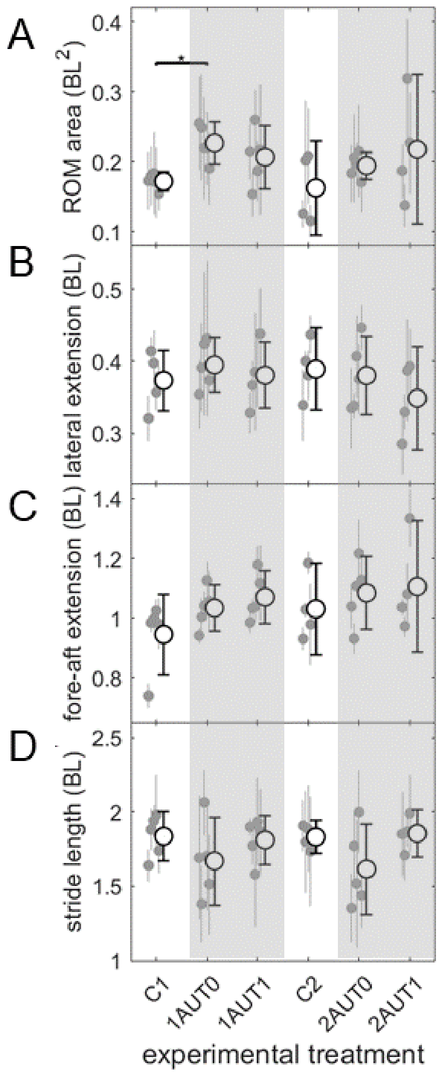

Table S1. The gait space metric tensor, $g_{ij}$, and maximum possible distance, $d_{max}$, between two points in gait space for locomotion with 6 and 8 legs.

| Number of legs | 6 | 8 |
|---|---|---|
| $g_{ij}$ | $\frac{1}{6}\begin{bmatrix} 5 & 4 & 3 & 2 & 1 \\ 4 & 8 & 6 & 4 & 2 \\ 3 & 6 & 9 & 6 & 3 \\ 2 & 4 & 6 & 8 & 4 \\ 1 & 2 & 3 & 4 & 5 \end{bmatrix}$ | $\frac{1}{8}\begin{bmatrix} 7 & 6 & 5 & 4 & 3 & 2 & 1 \\ 6 & 12 & 10 & 8 & 6 & 4 & 2 \\ 5 & 10 & 15 & 12 & 9 & 6 & 3 \\ 4 & 8 & 12 & 16 & 12 & 8 & 4 \\ 3 & 6 & 9 & 12 & 15 & 10 & 5 \\ 2 & 4 & 6 & 8 & 10 & 12 & 6 \\ 1 & 2 & 3 & 4 & 5 & 6 & 7 \end{bmatrix}$ |
| $d_{max}$ (cycles) | 2.09 | 3.24 |

Table S2. Results of Kruskal-Wallis tests comparing various measures for different treatments vs the C1 control.

The effect size is defined as the ratio between the ratio of the measure in question for the C1 control and the treatment (e.g. for speed, $v_{treatment}/v_{C1}$).

Speed

| Group | Rank Lower Limit | Rank Difference | Rank Upper Limit | P | Effect size |
|---|---|---|---|---|---|
| 1AUT0 | -36.1 | -20.1 | -4.10 | 0.01 | 0.72 |
| 1AUT1 | -24.7 | -8.7 | 7.30 | 0.57 | 0.86 |
| C2 | -18.9 | -1.8 | 15.33 | 1.00 | 0.94 |
| 2AUT0 | -35.1 | -19.1 | -3.10 | 0.01 | 0.74 |
| 2AUT1 | -21.2 | -4.0 | 13.08 | 0.98 | 0.92 |

Tortuosity

| Group | Rank Lower Limit | Rank Difference | Rank Upper Limit | P | Effect size |
|---|---|---|---|---|---|
| 1AUT0 | -25.4 | -9.5 | 6.53 | 0.48 | 0.99 |
| 1AUT1 | -18.4 | -2.5 | 13.53 | 1.00 | 1.00 |
| C2 | -12.0 | 5.1 | 22.26 | 0.95 | 1.00 |
| 2AUT0 | -20.0 | -4.1 | 11.93 | 0.98 | 0.99 |
| 2AUT1 | -14.2 | 2.9 | 20.01 | 1.00 | 1.00 |

Yaw

| Group | Rank Lower Limit | Rank Difference | Rank Upper Limit | P | Effect size |
|---|---|---|---|---|---|
| 1AUT0 | -39.6 | -23.7 | -7.67 | 0.001 | -9.0 |
| 1AUT1 | -34.0 | -18.1 | -2.07 | 0.019 | -6.9 |
| C2 | -27.5 | -10.4 | 6.76 | 0.452 | -4.1 |
| 2AUT0 | -34.6 | -18.7 | -2.67 | 0.014 | -7.6 |
| 2AUT1 | -39.0 | -21.9 | -4.74 | 0.005 | -8.8 |

Stride frequency

| Group | Rank Lower Limit | Rank Difference | Rank Upper Limit | P | Effect size |
|---|---|---|---|---|---|
| 1AUT0 | -28.9 | -15.8 | -2.70 | 0.011 | 0.67 |
| 1AUT1 | -21.5 | -8.4 | 4.70 | 0.355 | 0.82 |
| C2 | -16.8 | -2.9 | 10.99 | 0.980 | 0.92 |
| 2AUT0 | -26.7 | -13.6 | -0.50 | 0.039 | 0.74 |
| 2AUT1 | -19.0 | -5.2 | 8.74 | 0.823 | 0.93 |

Duty factor

| Group | Rank Lower Limit | Rank Difference | Rank Upper Limit | P | Effect size |
|---|---|---|---|---|---|
| 1AUT0 | 3.3 | 16.4 | 29.50 | 0.007 | 1.07 |
| 1AUT1 | -2.5 | 10.6 | 23.70 | 0.158 | 1.04 |
| C2 | -4.4 | 9.5 | 23.34 | 0.300 | 1.03 |
| 2AUT0 | 3.3 | 16.4 | 29.50 | 0.008 | 1.07 |
| 2AUT1 | -2.7 | 11.2 | 25.09 | 0.161 | 1.06 |

Static stability margin (SSM)

| Group | Rank Lower Limit | Rank Difference | Rank Upper Limit | P | Effect size |
|---|---|---|---|---|---|
| 1AUT0 | -27.1 | -14.0 | -0.90 | 0.031 | 0.41 |
| 1AUT1 | -28.9 | -15.8 | -2.70 | 0.011 | 0.39 |
| C2 | -14.8 | -0.9 | 12.99 | 1.000 | 0.87 |
| 2AUT0 | -23.9 | -10.8 | 2.30 | 0.146 | 0.60 |
| 2AUT1 | -31.5 | -17.7 | -3.76 | 0.006 | 0.37 |

Foot ROM area

| Group | Rank Lower Limit | Rank Difference | Rank Upper Limit | P | Effect size |
|---|---|---|---|---|---|
| 1AUT0 | 1.3 | 14.4 | 27.50 | 0.02 | 1.3 |
| 1AUT1 | -4.1 | 9.0 | 22.10 | 0.29 | 1.2 |
| C2 | -12.2 | 1.7 | 15.54 | 1.00 | 1.0 |
| 2AUT0 | -6.7 | 6.4 | 19.50 | 0.62 | 1.1 |
| 2AUT1 | -4.5 | 9.4 | 23.29 | 0.30 | 1.2 |

Lateral leg extension

| Group | Rank Lower Limit | Rank Difference | Rank Upper Limit | P | Effect size |
|---|---|---|---|---|---|
| 1AUT0 | -9.3 | 3.8 | 16.90 | 0.925 | 1.06 |
| 1AUT1 | -12.3 | 0.8 | 13.90 | 1.000 | 1.02 |
| C2 | -10.3 | 3.6 | 17.49 | 0.952 | 1.01 |
| 2AUT0 | -11.9 | 1.2 | 14.30 | 1.000 | 1.02 |
| 2AUT1 | -17.0 | -3.2 | 10.74 | 0.972 | 0.99 |

Fore-after leg extension

| Group | Rank Lower Limit | Rank Difference | Rank Upper Limit | P | Effect size |
|---|---|---|---|---|---|
| 1AUT0 | -5.7 | 7.4 | 20.50 | 0.480 | 1.09 |

| | | | | | |
|---|---|---|---|---|---|
| 1AUT1 | -3.5 | 9.6 | 22.70 | 0.234 | 1.13 |
| C2 | -9.7 | 4.2 | 18.04 | 0.916 | 1.08 |
| 2AUT0 | -2.3 | 10.8 | 23.90 | 0.146 | 1.15 |
| 2AUT1 | -4.5 | 9.4 | 23.29 | 0.305 | 1.17 |

Stride length

| Group | Rank Lower Limit | Rank Difference | Rank Upper Limit | P | Effect size |
|---|---|---|---|---|---|
| 1AUT0 | -20.3 | -7.2 | 5.90 | 0.507 | 0.91 |
| 1AUT1 | -14.1 | -1.0 | 12.10 | 1.000 | 0.99 |
| C2 | -14.3 | -0.4 | 13.49 | 1.000 | 1.01 |
| 2AUT0 | -20.7 | -7.6 | 5.50 | 0.454 | 0.88 |
| 2AUT1 | -14.0 | -0.1 | 13.74 | 1.000 | 0.96 |

Table S3. Results of Wilcoxon rank sum tests comparing various measures between two treatments.

The effect size is defined as the ratio between the ratio of the measure in question for the first and second treatment (e.g., for speed, $v_{2AUT0}/v_{1AUT0}$). NC = not computed.

| Treatments compared | 2AUT0 & 1AUT0 | | 1AUT0 & CC | |
|---|---|---|---|---|
| Variable | P | Effect size | P | Effect size |
| speed | 1.00 | 1.04 | 0.476 | 0.96 |
| tortuosity | 0.55 | 1.01 | 0.914 | 0.99 |
| yaw | 0.22 | 0.85 | NC | NC |
| Stride frequency | 0.69 | 1.10 | NC | NC |
| Duty factor | 1.00 | 1.00 | NC | NC |
| SSM | 0.31 | 1.45 | NC | NC |
